# YAP1 drives aggressive and therapy resistant state in melanoma through reprogramming the chromatin and regulating immune evasive programs

**DOI:** 10.1101/2025.09.10.675381

**Authors:** Nana Chen, Gracia Bonilla, Abdullah AI Emran, Brian Lin, Haymanti Bhanot, Yang Sun, Avinash Waghray, Mateo Useche, David Liu, Genevieve Boland, Ivana de la Serna, Valentina Sancisi, Martin Sattler, Jayaraj Rajagopal, Xu Wu, Ruslan I. Sadreyev, David Fisher, Srinivas Vinod Saladi

## Abstract

Despite promising initial results in targeting the RAF-MEK-ERK cascade, resistance to BRAF/MEK inhibitors remains a critical challenge in nearly 50% of melanoma patients. Our study demonstrates that robust YAP1 activation in metastatic melanoma correlates with poor survival and drives transcriptional programs linked to therapeutic resistance. Mechanistically, YAP1 predominantly remodels the chromatin landscape in resistant tumors by partnering with BRD4 and TEAD, creating a permissive transcriptional state that sustains oncogenic signaling. Clinical validation in biopsies from resistant melanoma confirms elevated expression of YAP1 target genes. Furthermore, pharmacological inhibition of BRD4 or TEAD reduces YAP1-driven transcription and reactivates antitumor immunity programs. TEAD specific inhibitors (and not verteporfin which is a highly non-specific inhibitor) synergize with immune checkpoint blockade in in vivo model system by promoting increased CD8⁺ T cell infiltration and prolonged survival in the melanoma mouse model. Collectively, these findings reveal a chromatin-centric vulnerability in BRAF/MEK inhibitor-resistant melanoma and propose TEAD specific inhibitors as a promising dual strategy to overcome resistance and reinvigorate the immune response, offering a novel therapeutic avenue for patients.

Resistance to BRAF and MEK inhibitors remains a major obstacle in the treatment of melanoma. Here, we show that elevated YAP1 activity in metastatic melanoma is associated with poor patient survival and drives transcriptional programs that promote therapeutic resistance. Mechanistically, YAP1 cooperates with BRD4 and TEAD to reprogram the chromatin landscape, establishing an oncogenic transcriptional state that sustains resistance. Analysis of patient-derived melanoma biopsies confirms increased expression of YAP1 target genes in resistant tumors. Targeting this epigenetic circuitry using pharmacological inhibitors of BRD4 or TEAD effectively suppresses YAP1-driven transcriptional output and restores antitumor immune activity, as evidenced by enhanced CD8⁺ T cell infiltration and prolonged survival in resistant melanoma models. These effects are further supported by transcriptomic and chromatin accessibility analyses, which reveal reduced expression of immune-suppressive and proliferation-associated gene networks upon TEAD inhibition. Collectively, our findings identify a chromatin-centric mechanism underlying resistance to MAPK-targeted therapy and nominate TEAD inhibition as a promising dual-action strategy to both overcome resistance and re-engage antitumor immunity. This work offers a compelling therapeutic avenue for patients with drug-resistant melanoma.

## 1. Introduction

Genetic and environmental factors contribute to the malignant transformation of melanocytes into melanomas ^[1]^. Approximately half the patients with metastatic melanoma have a BRAF mutation, which, along with RAS mutations, leads to oncogenic activation of mitogen-activated protein kinase (MAPK) and other signaling pathways to promote tumor cell survival ^[2]^. However targeting the MAPK pathway with RAF and MEK inhibitors has shown limited clinical efficacy in melanomas with BRAF V600E mutation, due to primary and acquired resistance ^[3]^. Therefore, identifying new molecular targets and therapeutic strategies to limit this resistance is of great interest.

Many factors contribute to resistance to BRAF inhibitors (BRAFi) ^[2b]^. One of these is actin remodeling, which can lead to YAP1/TAZ accumulation in the nucleus, thereby promoting cell survival and proliferation ^[4]^. The YAP1/TAZ/TEAD complex has been shown to regulate expression of the anti-apoptosis gene BCL-XL to protect BRAF– and RAS-mutated tumor cells from death ^[2c]^. Indeed, BRAF inhibitor-resistant melanoma cancer stem cells express high levels of YAP1, TAZ, and TEAD ^[5]^. Thus, YAP1/TAZ/TEAD and their interacting partners including epigenetic factors are promising and important drug targets in BRAFi-resistant melanoma. Recently, inhibitors targeting the interaction between YAP1 and TEAD have been developed, including Verteporfin^[6]^ and super-TDU^[7]^. However, these early compounds exhibit relatively low specificity and limited efficacy, and Verteporfin is a highly non-specific compound targeting multiple pathways ^[8]^. To overcome this, more recently, a new generation of selective TEAD inhibitors has emerged, such as IK-930 ^[9]^, K-975 ^[10]^, VT3989^[11]^, and MGH-CP1^[12]^, which block TEAD auto-palmitoylation and have demonstrated improved potency and selectivity in preclinical models. Several of these compounds have advanced into early-phase clinical trials, reflecting the growing therapeutic interest in targeting the YAP/TEAD axis in cancers with Hippo pathway alterations or transcriptional addiction to YAP/TAZ activity. These advances underscore the urgent need for highly specific and efficacious inhibitors that can disrupt YAP1 or its critical interactors in a clinically meaningful way.

As a key Hippo pathway effector, YAP1 has emerged as a critical immune regulator in tumor cells ^[13]^. Activation of YAP1 is involved in the suppression of anti-tumor immune responses ^[14]^. In BRAFi-resistant melanoma, YAP1 upregulates PD-L1 (CD274) to mediate the evasion of cytotoxic T-cell mediated tumor killing^[15]^. Moreover, the Hippo-YAP1 signaling pathway regulates polarization of tumor-associated macrophages (TAMs) to promote tumor progression ^[16]^. Finally, an activated YAP1 signature is associated with the expression of myeloid-derived suppressor cell (MDSC) related genes and drives MDSC recruitment ^[17]^. Collectively, these findings position YAP1 as a critical modulator of immunosuppression in melanoma, and suggest that targeting YAP1 could restore antitumor immunity and improve therapeutic outcomes.

Aberrant activation of transcriptional programs mediated by epigenome deregulation plays a critical role in melanoma pathogenesis ^[18]^. Specifically, bromodomain-containing protein 4 (BRD4) is reported to be a coactivator of YAP1 and is responsible for activating transcriptional programs ^[19]^. BRD4 is a member of the BET protein family enriched at promoters and enhancers, where it interacts with transcription factors to promote the expression of lineage-specific genes ^[20]^. BRD4 is a chromatin reader and writer that mainly recognizes and maintains lysine acetylation at H3 and H4 histones or transcriptional regulators at DNA regulatory elements ^[21]^. Importantly, YAP1/TAZ/TEAD/BRD4 interactions and co-localization onto promoter or enhancer regions are required for maintaining high transcription levels of genes that contribute to oncogenic transcriptional programs ^[19, 22]^. Recently, we demonstrated that FAT1-mutated HNSCC cells exhibit an oncogenic chromatin state (OCS) mediated by the YAP1/TAZ/TEAD transcription complex recruiting BRD4 and promoting H3K27 acetylation, and oncogenic transcriptional program ^[23]^. These studies suggest that transcriptional or epigenetic coregulators, including YAP1 or BRD4, are attractive therapeutic targets for treating cancer by overcoming resistance to current first-line treatments through targeting the OCS.

In this study, we investigated the role of YAP1 and its interactors in BRAF– and MEK inhibitor resistant melanoma and focused on BRAF-mutated melanomas that developed resistance to BRAFi, Vemurafenib or Dabrafenib and MEKi, Trametinib after chronic exposure. Specifically, we utilized comprehensive transcriptomic and epigenetic analyses to investigate the role of the Hippo pathway in promoting a treatment resistant state in melanoma. We also reveal that treatment with the BRD4 inhibitor JQ1 or the novel TEAD inhibitor MGH-CP1 ^[24]^ can abrogate the transcriptional activity of YAP1 and improve the immune response in MEK inhibitor-resistant melanoma cells.

## 2. Results

### 2.1 YAP1 and its target genes are highly enriched in metastatic melanoma and correlate with poorer prognosis in patients

The activating mutation of BRAF is well established as a key oncogenic driver in melanomagenesis and has also been implicated in facilitating tumor progression and metastatic dissemination^[25]^. Given the emerging role of Hippo-YAP1 signaling in therapy adaptation, we systematically analyzed its genomic and transcriptomic alterations in melanoma progression. First, we queried the TCGA data set for melanoma patients to see if Hippo pathway components are altered. Comprehensive genomic profiling of Hippo pathway components revealed recurrent genetic alterations indicative of pathway dysregulation. FAT4 and FAT1, key upstream regulators, exhibited the highest overall alteration frequencies, predominantly through loss-of-function mutations. Intermediate components such as TAOK2, MST2 (STK3), and LATS1 also showed notable frequencies of mutation and deep deletion (**Figure 1A**). In contrast, the core downstream effectors YAP1 and WWTR1 (TAZ) were primarily affected by gene amplification, consistent with their established oncogenic roles (**Figure 1A**). Transcriptomic analyses demonstrated significantly elevated expression of YAP1, WWTR1, TEAD3, and TEAD4 in metastatic versus primary melanoma (**Figures 1B and S1A**). To further investigate the prognostic relevance of Hippo-YAP/TAZ pathway components and their associated gene expression in human cancers, we performed survival analyses using the TIDE platform^[26]^ across multiple public datasets, including TCGA and immunotherapy-treated cohorts. Kaplan-Meier survival curves revealed that high expression of YAP1 protein levels, along with elevated WWTR1, TEAD3, and TEAD4 mRNA expression and their target genes, were associated with reduced survival probability in both primary and metastatic melanoma **(Figures 1C and S1B)**. In metastatic melanoma, YAP1 expression exhibited a significant positive correlation with expression of its target genes, including *CTGF*, *CYR61*, *AXL*, *BCL2L1*, *FSTL1*, *NRG1*, *ANKRD1*, *AMOLT1*, and *FST* (**Figures 1D and S1C**). Moreover, *CTGF*, *CYR61*, *AXL*, *BCL*-*XL*, and *FSTL1* showed significantly higher expression in metastatic melanoma than in primary melanoma (**Figure 1E**). Notably, AXL, and BCL-XL are established mediators of targeted therapy resistance in solid tumors^[27]^. Single-cell RNA-seq analysis revealed elevated expression of YAP1 and its downstream targets (*CTGF*, *CYR61*, *AXL*, *BCL-XL*, and *FSTL1*) within distinct subpopulations across multiple melanoma samples, highlighting intratumoral transcriptional heterogeneity and the selective activation of the YAP1 transcriptional program in melanoma (**Figure 1F**) ^[28]^. Collectively, these findings position YAP1 activation as a convergence point for melanoma metastasis and therapy resistance.

**Figure 1.**
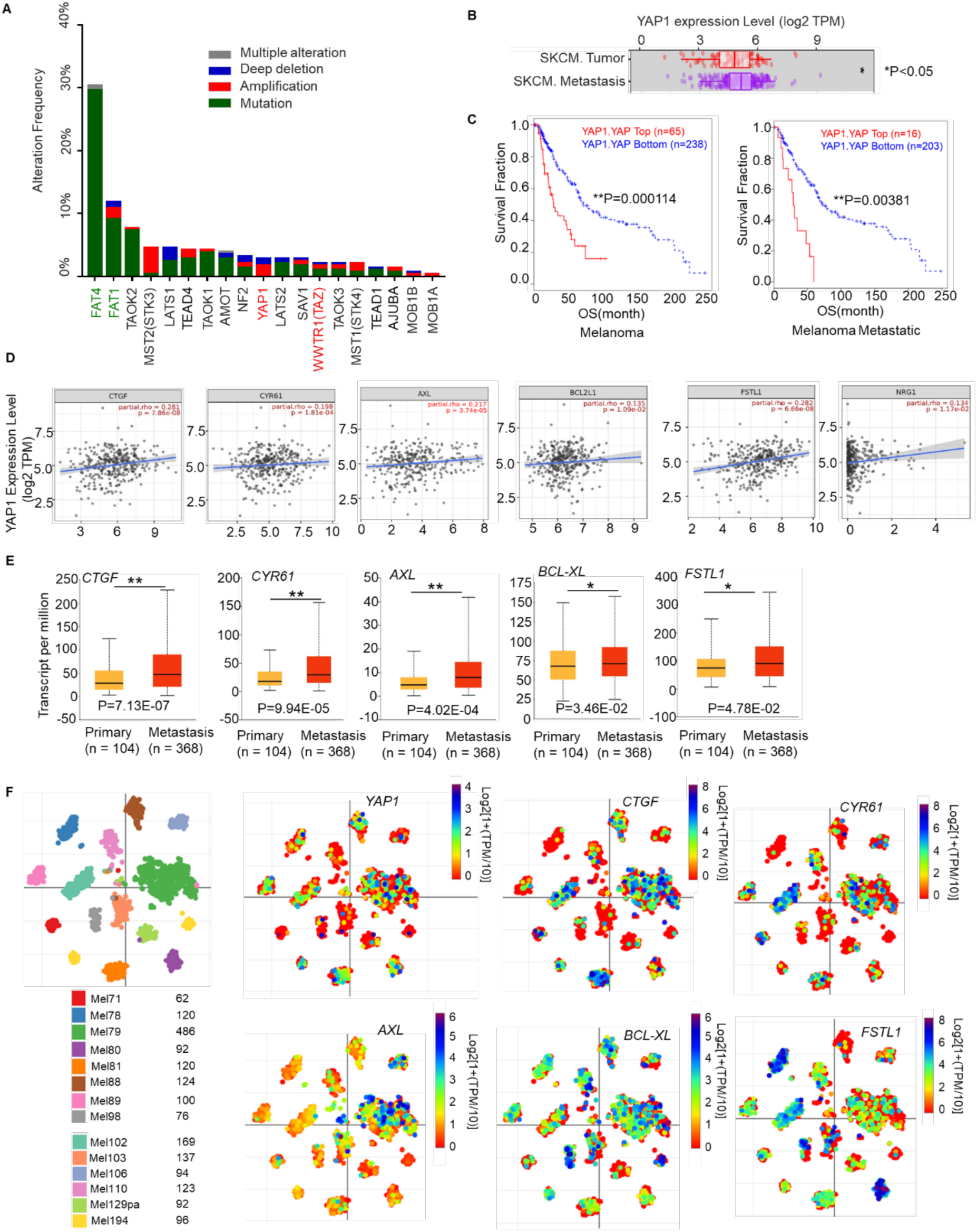
YAP1 and its target genes are highly expressed in metastatic melanoma and correlated with poor survival. A) The alteration frequency of Hippo pathway core genes in SKCM from TCGA. The genes with high somatic alteration frequency are highlighted in green, YAP1 and WWTR1 were highlighted in red. B) The expression level of YAP1 in SKCM primary and metastatic tumors. (*P<0.05). C) Correlation between YAP1 protein expression and survival fraction in primary and metastatic melanoma. D) The correlation between YAP1 and its target genes in metastatic melanoma (adjusted tumor purity). E) The expression level of YAP1 target genes *CTGF*, *CYR61*, *AXL*, *BCL-XL*, and *FSTL1*, in primary and metastasis melanoma. (* P<0.05; **P<0.01). F) Single cell RNA-seq analysis of *YAP1* and its target genes in melanoma patients. tSNE plot of malignant cells (dots) from clinical melanoma tumors. Only tumors with at least 50 malignant cells are shown.

### 2.2 Transcriptomic and chromatin accessibility profiling reveal YAP1 activation-driven therapy resistance in melanoma

To directly interrogate whether YAP1 activation is linked to therapeutic resistance in melanoma patients, we analyzed transcriptomic profiles from BRAF V600E metastatic melanoma patients who developed resistance following BRAF inhibitor (dabrafenib) and MEK inhibitor (trametinib) combination therapy (GSE77940)^[29]^ (**Figure 2A**). We specifically investigated whether YAP1 target genes^[30]^ were implicated in driving this resistance by examining gene expression data from pre– and post-BRAF/MEK inhibitor treatment biopsies in these patients. We found robust upregulation of multiple YAP1 target genes, including *CTGF*, *CYR61*, *AXL*, *BCL2L1*, *NRG1*, *ANKRD1*, *FST*, and *CD274* in resistant tumors **(Figure 2B)**. These findings suggest that YAP1-driven transcriptional reprogramming is a key feature of the acquired resistance phenotype and may play a central role in mediating the adaptive response to MAPK-targeted therapy in melanoma. To mechanistically dissect how Hippo-YAP1 signaling promotes therapy resistance, we performed ATAC-seq in isogenic MEKi-sensitive and MEKi-resistant A375 melanoma cells^[31]^. Analysis of the ATAC-seq data revealed striking differences with increased chromatin accessibility at YAP1/TAZ target genes, including *CYR61*, *AXL*, *BCL*-*XL*, *NRG1*, and *CD274* in MEKi resistant A375 cells (**Figure 2C**), suggesting epigenetic reprogramming as a potential mechanism driving resistance. Motif enrichment analysis revealed a shift in transcriptional regulation underlying MEKi resistance. Parental A375 cells exhibited strong SOX10, SOX6, and SOX2 motifs, indicating active SOX-driven transcriptional programs that maintain melanoma lineage identity. Additionally, enrichment of BATF, FOS, and FOSB motifs suggests active AP-1 signaling, which is linked to stress responses and adaptive transcription. The presence of TFAP2 family (TFAP2A, TFAP2B, TFAP2C) and ZNF24 motifs implies roles in differentiation and chromatin regulation. In contrast, MEKi-resistant cells showed notable loss of SOX motifs was detected, indicating lineage dedifferentiation. Concurrently, there was a higher enrichment of TEAD1, TEAD3, and TEAD4 motifs, suggesting activation of the YAP/TEAD axis, which is known to drive tumor plasticity and drug resistance. These findings illustrate a shift from SOX/AP-1-driven transcription to TEAD/YAP-mediated regulation, highlighting the YAP/TEAD axis as a key factor in the transcriptional reprogramming in MEKi-resistant melanoma cells (**Figure 2D**). To investigate how changes in chromatin accessibility influence functional pathway regulation in MEKi-resistant versus parental A375 cells, we performed KEGG pathway enrichment analysis. Regions with increased chromatin accessibility in resistant cells were significantly associated with genes that are components of the Hippo signaling pathway, consistent with motif enrichment analysis suggesting TEAD activation (**Figure 2E**). To validate these findings, we used qPCR with specific primers to assess chromatin accessibility at multiple Hippo pathway target genes. Our analysis confirmed increased chromatin accessibility at the promoters of YAP1 target genes (*CYR61*, *AXL*, *BCL-XL*) as well as immune-related gene (*CD274*) in MEKi-resistant A375 cells compared to parental A375 cells (**Figure S2A**). Together, these findings support that upregulation of Hippo-YAP1 signaling contributes to the development of MEKi resistance in melanoma.

**Figure 2.**
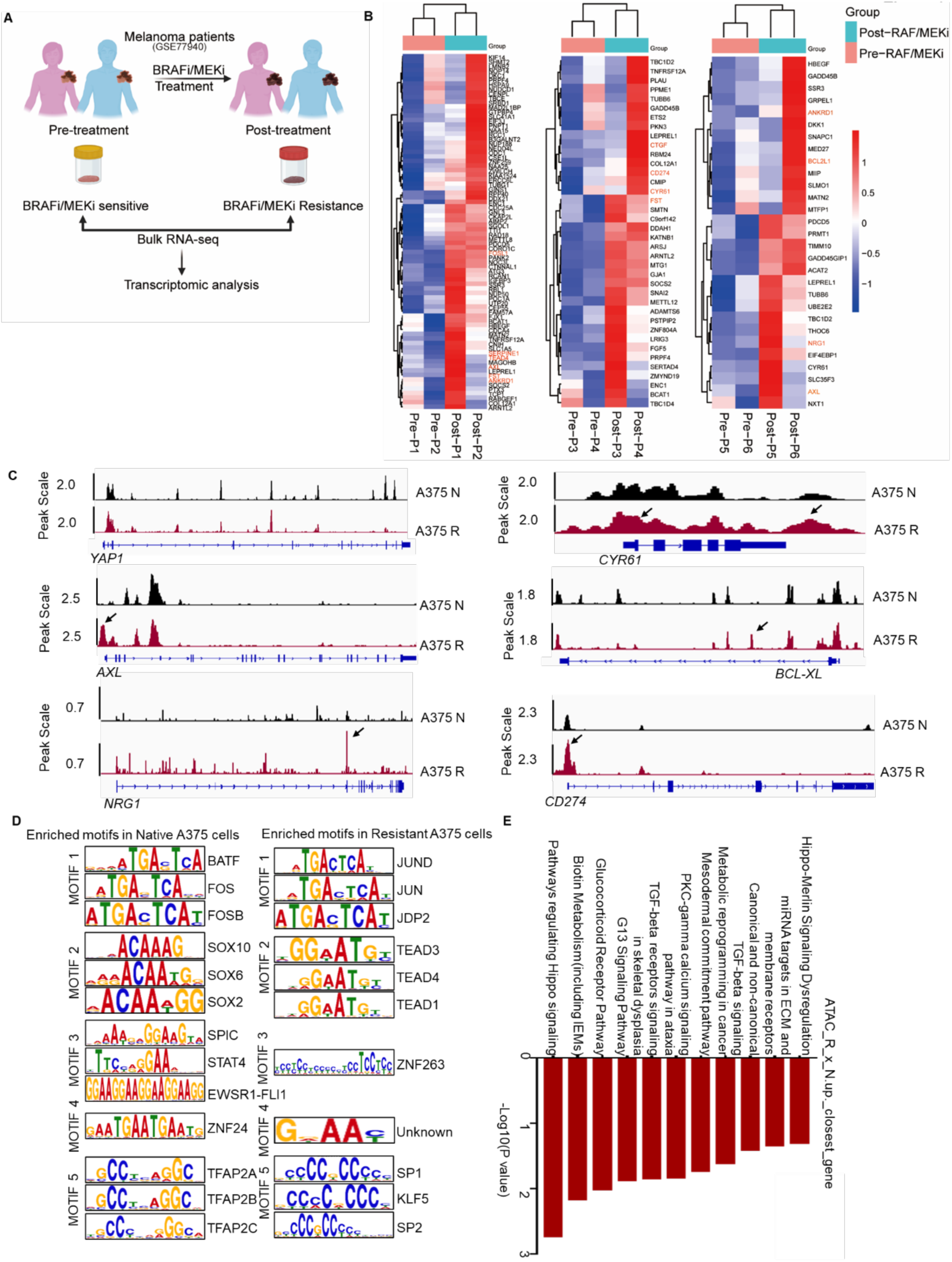
Transcriptomic and chromatin accessibility profiling reveal YAP1 activation-driven therapy resistance in melanoma. A) Schematic representation of transcriptomic analysis workflow for melanoma patient samples before and after BRAFi/MEKi treatment (BRAFi/MEKi resistant) patient samples from the GSE77940 dataset. B) Heatmap depicting changes in expression of YAP1 target genes between pre-treatment (Pre) and post-treatment resistant (Post) melanoma patient samples from the GSE77940 dataset. Genes highlighted in red represent key YAP1 targets identified in this study. C) Genomic tracks of ATAC-seq (normalized) signal at *YAP1, CYR61, AXL, BCL2L1, NRG1,* and *CD274* locus in A375 Native and A375 Resistant cells. D) HOMER motif enrichment analyses of differentially accessible peaks (A375 R versus A375 N). The enrichment of binding motifs of transcription factors per sample are shown. E) KEGG pathway analyses showed distinct pathways linked to differentially accessible peaks (A375 R versus A375 N).

### 2.3 YAP1 transcriptional activation is involved in MEK inhibitor resistance

To further elucidate the mechanistic role of YAP1 to therapeutic resistance, we performed RNA-seq comparing parental and MEKi-resistant melanoma cells **(Figure S3A)**. We found that well established YAP1 target genes including *CTGF*, *CYR61*, *ANKRD1*, *SERPINE1*, *FSTL1*, and *FOSL1* were up-regulated in resistant cells, reinforcing YAP1 activation as a key transcriptional driver **(Figure 3A)**. Additionally, proliferation associated YAP1 target genes, including *AXL*, *NRG1*, *EGFR*, and *EPHA2*, as well as anti-apoptosis YAP1 target genes (*BCL*-*XL* and *BIRC3*), were significantly upregulated, suggesting enhanced survival and growth potential in resistant cells **(Figure 3A)**. This was accompanied by increased expression of genes involved in invasion (*MMP1*, *MMP3*, *MMP10*, and *MMP12*), indicating a shift toward a more aggressive phenotype **(Figure 3B)**. In contrast, melanocytic lineage genes such as *MITF*, *SOX10*, *TYR*, *PMEL*, and *DCT* were downregulated, consistent with dedifferentiation and loss of melanoma identity **(Figure 3B)**. Furthermore, resistant cells exhibited immune evasion features, with downregulation of anti-tumor immune response genes (*IL-24*, *CD74*, and *HLA-DRA*) and upregulation of *CD274* **(Figure 3B)**, indicating that the resistant cells exhibit an immune suppressive phenotype. To compare the global YAP1 dependent transcriptome in resistant versus parental cells, we performed RNA-seq following knock down of YAP1 in both A375 parental and A375 MEKi resistant cells. YAP1 knockdown consistently downregulated YAP1 target genes and up-regulated immune response genes in A375 MEKi resistant cells, while no such effects were observed in A375 parental cells **(Figure 3C)**. Furthermore, KEGG pathway analysis for genes downregulated upon YAP1 knockdown in MEKi resistant cells revealed enrichment in the Hippo signaling and stem cell signaling pathways **(Figure 3D)**. Conversely, upregulated genes following knock down of YAP1 in resistant cells were associated with the defense response to viruses, cytokine-mediated signaling pathways, and cytokine-cytokine receptor interaction **(Figure 3E)**, suggesting that YAP1 is a key effector mediating MEK inhibitor resistance and immune suppression. Moreover, YAP1 knockdown in MEK inhibitor-resistant cells led to a significant reduction in the expression of YAP1 target genes (*CTGF*, *CYR61*, *ANKRD1*, *NRG1*). Additionally, we observed a decrease in *CD274* expression **(Figure 3F)**. Collectively, these findings indicate that YAP1 contributes to a proliferative, metastatic, and immune-suppressive expression signatures in MEK inhibitor-resistant cells.

**Figure 3.**
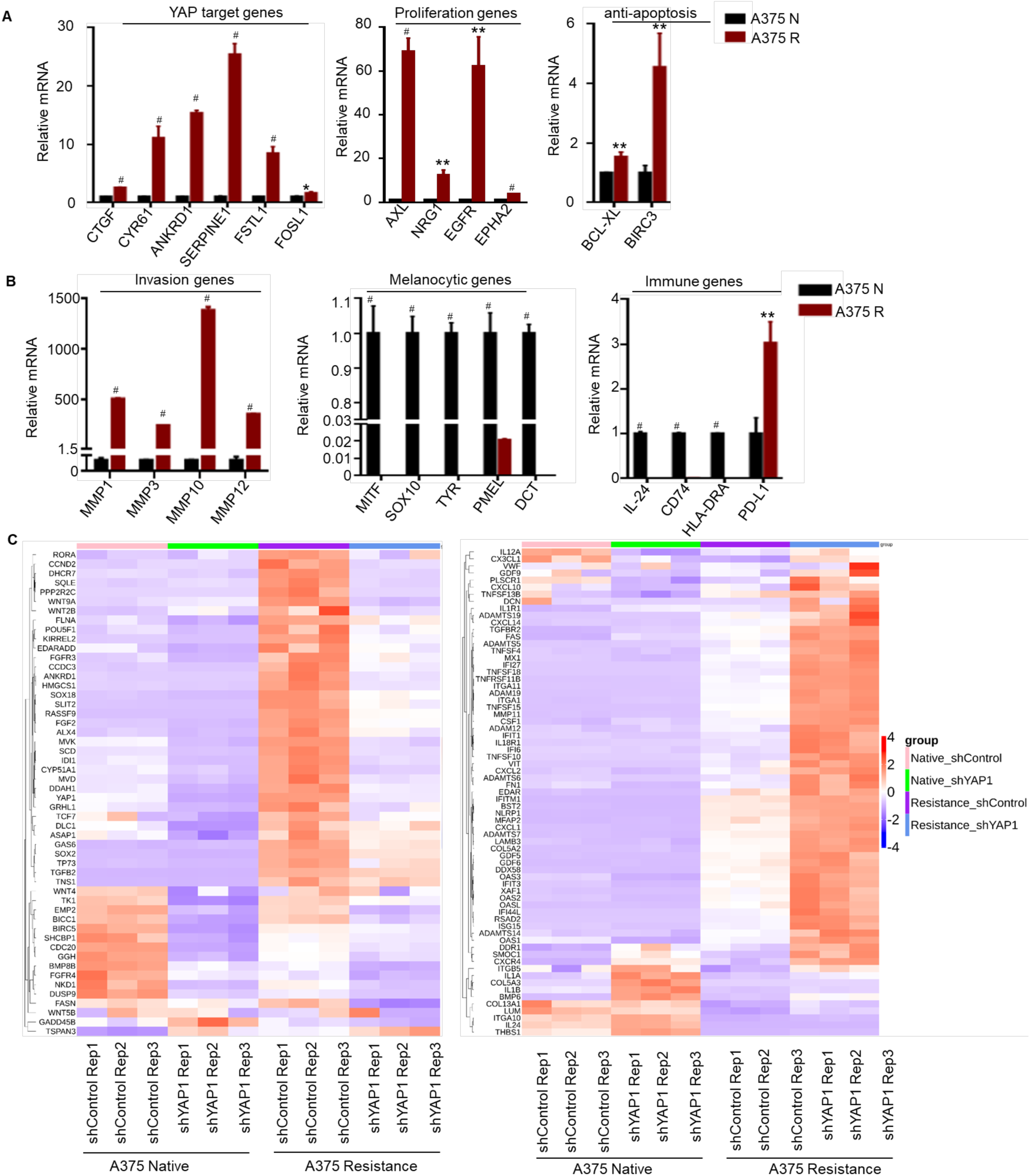
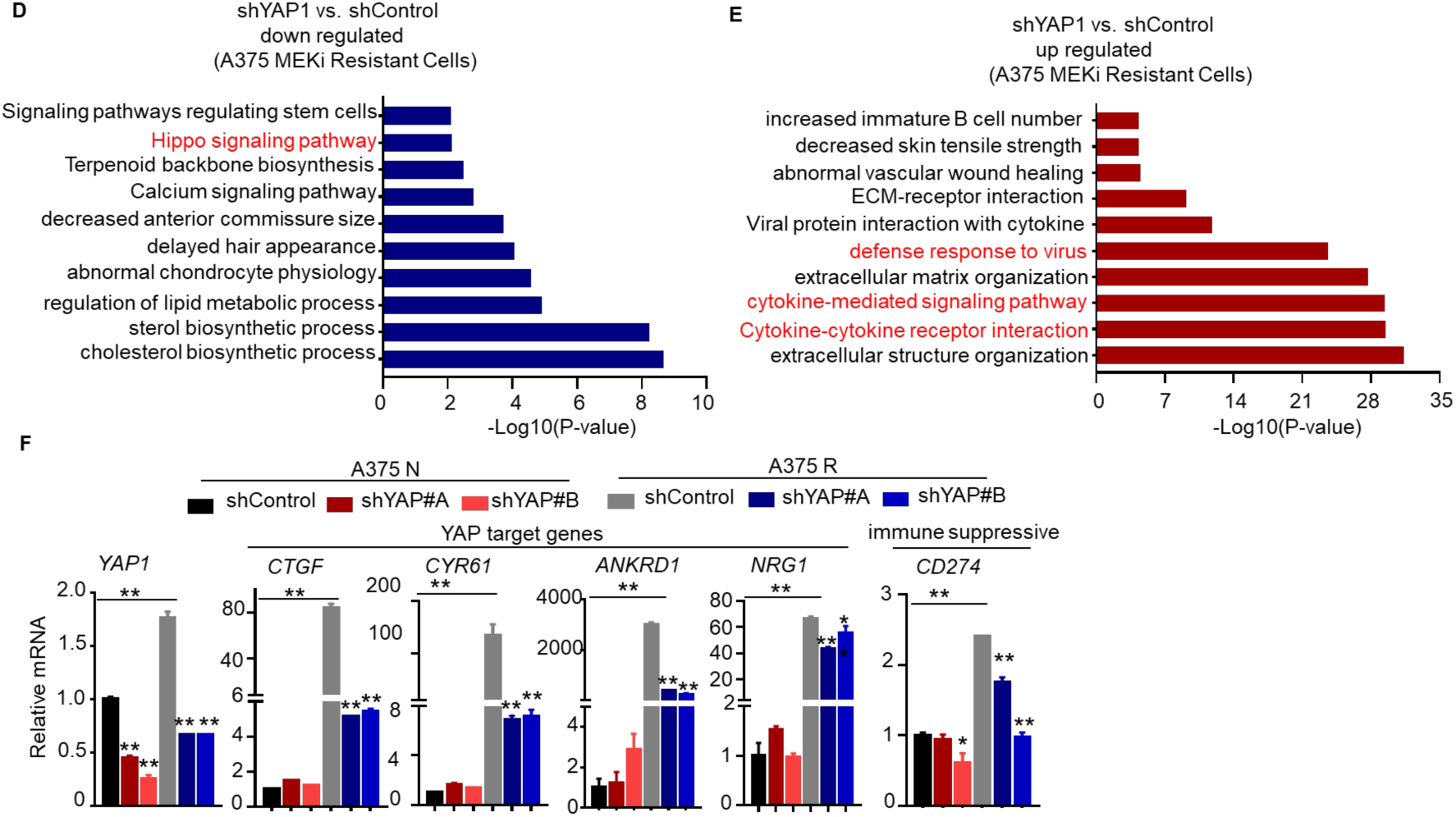
Transcriptional activation of YAP1 is involved in resistance to MEK inhibitor and immune suppression. A, B) Relative expression of indicated genes in A375 Native and A375 Resistant cells by qPCR analysis (n = 3 independent experiments, graph shows mean ± SD, *p<0.05, **p<0.01, ^#^p<0.001). C) Heatmaps for normalized RNA-Seq data (scaled and centered) from A375 native and MEKi resistant cells after YAP1 knock down. Heatmap shows the YAP1 regulated genes (Left) and immune related genes (Right). D, E) KEGG pathway enrichment analyses of DEGs by RNA-seq upon YAP knockdown in A375 Resistant cells. The x-axis indicates the –log10 (p-value). The y-axis indicates the names of KEGG pathways. F) Quantification of YAP1 target genes and immune genes with knockdown of *YAP1* in A375 Native and A375 Resistant cells (*n* = 3 independent experiments, error bar shows mean ± SD, *P<0.05, **P<0.01).

### 2.4 YAP1 cooperates with BRD4 to regulate its target genes by promoting changes in chromatin accessibility

To identify potential interactors of YAP1, we queried the BioGRID database^[32]^, integrating published data ^[19, 23, 33]^, and gene sets from MEKi resistant cells ^[34]^. We found that BRD4 was enriched as an interacting partner of YAP1 **(Figure S4A)**. We further conducted Chromatin Immunoprecipitation Enrichment Analysis (ChEA) to identify transcription factors and cofactors that may regulate genes driving the MEK inhibitor-resistant state. This analysis revealed BRD4 as a key regulator for a majority of resistance-associated genes, suggesting that BRD4 cooperates with YAP1 to promote MEK inhibitor resistance **(Figure S4B)**. To investigate whether these genes were directly regulated by YAP1 and BRD4, we performed Chromatin Immunoprecipitation (ChIP) studies for YAP1, WWTR1, TEAD4, and BRD4 in both parental and MEK inhibitor-resistant cell lines. Notably, YAP1 and BRD4 exhibited significant co-enrichment in the promoter regions of *NRG1*, *AXL*, *PD*-*L1*, and *BCL*-*XL* in MEKi resistant cells **(Figure 4A)**. Conversely, TAZ and TEAD4 displayed robust enrichment only in the promoter regions of *AXL* and *BCL*-*XL*, indicating that YAP1 and BRD4 co-occupy the promoter regions of Hippo-YAP targeted genes in the context of MEK inhibitor resistance **(Figure 4A)**. Furthermore, there was a significant increase in the enrichment of H3K27ac at the loci of YAP1 binding sites on its target genes in MEKi resistant cells, suggesting activation of their promoter or enhancer regions **(Figure 4B)**. Taken together, these findings strongly suggest a cooperative regulatory role for YAP1 and BRD4 in the activation of Hippo-YAP1 target genes in the context of MEK inhibitor resistance.

**Figure 4.**
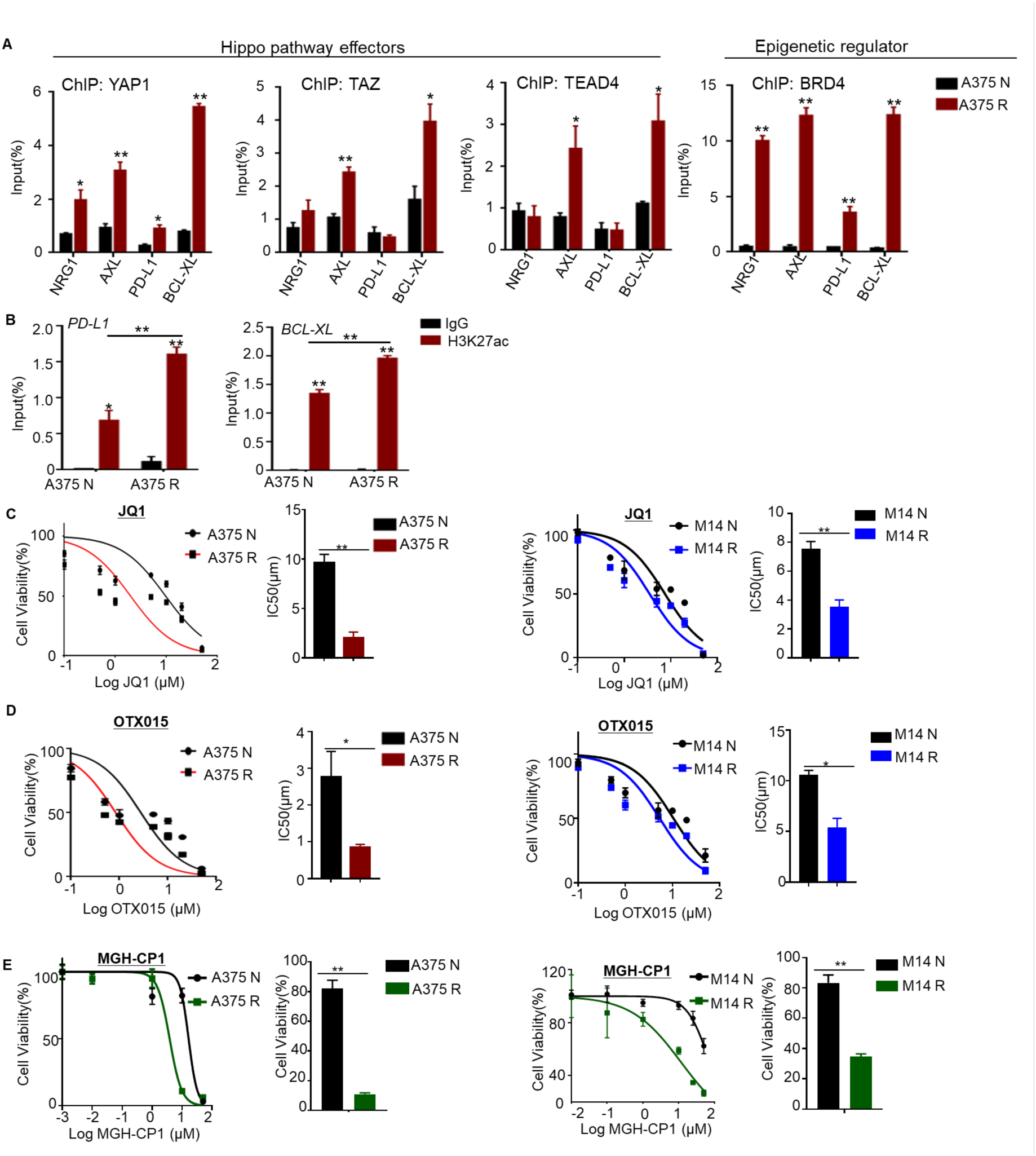
BRAFi/MEKi resistant melanoma cells are sensitive to YAP co-effectors inhibitors. A) Analysis of YAP1, TAZ, TEAD4 and BRD4 binding to *NRG1*, *AXL*, *PD-L1* and *BCL-XL* promoter loci in A375 N and A375 R cells by ChIP-qPCR. ChIP with pre-immune IgG displayed background signal (which was comparable in all samples). DNA enrichment was calculated and presented as fraction of input (n=3, *P<0.05, **P<0.01). B) ChIP-qPCR analyzing H3K27ac binding to promoters of *PD-L1* and *BCL-XL* in A375 N and A375 R cells. ChIP with pre-immune IgG displayed background signal (which was comparable in all samples). DNA enrichment was calculated as fraction of input and is presented as % of H3K27ac binding (n=3, *P<0.05, **P<0.01). C, D) Dose-response analysis of JQ1 and OTX015 in A375 N and A375 R, M14 N and M14 R. Data are mean of n=3 independent wells (independently treated and evaluated). Bar graph indicates the value of IC50 (n=3, *P<0.05, **P<0.01). E) Dose-response curve of A375 N, A375 R, M14 N and M14 R cells after 72h treatment with MGH-CP1. Each point shows mean ± S.D after normalization to DMSO-treated cells (n =3, **P<0.01).

### 2.5 MEK inhibitor resistant melanoma cells are selectively sensitive to BRD4 and TEAD inhibitors

It has been previously demonstrated that YAP/TAZ-TEAD transcriptional activity relies on the presence of BRD4 for genome-wide chromatin association ^[19]^. This interaction was sensitive to BET inhibitor JQ1, which abolished the transcriptional effect of YAP-TAZ in a breast cancer cell line^[19]^ and in HNSCC^[23]^. Additionally, BRD4 is significantly upregulated in metastatic melanoma, and is associated with poor survival **(Figures S4C-S4D)**. To explore whether the YAP1 transcriptional program could be therapeutically targeted through BRD4, we tested BRD4 inhibitors in MEKi-resistant cells. As hypothesized, resistant cells showed increased sensitivity to JQ1 and OTX015 compared to parental controls **(Figures 4C-4D)**, indicating a functional dependency on BRD4. Since TEADs are core mediators of YAP/TAZ signaling, we further evaluated MGH-CP1, a selective TEAD auto-palmitoylation inhibitor that disrupts YAP-TEAD interaction and downstream transcription ^[35]^. Consistent with our hypothesis, MEKi– and BRAFi-resistant cells exhibited greater sensitivity to MGH-CP1 than parental cells **(Figure 4E)**, suggesting that therapeutic resistance is associated with increased dependency on YAP-TEAD driven transcription. Together, these findings indicate that BRAFi/MEKi-resistant cells are particularly reliant on the YAP1-TEAD-BRD4 interaction to sustain their resistant phenotype and may be vulnerable to disruption of this transcriptional network.

### 2.6 Inhibition of BRD4 and TEAD affects the global chromatin landscape in MEK inhibitor resistant cells

To further elucidate the epigenetic mechanisms underlying BRAFi/MEKi resistance and the heightened sensitivity of MEK inhibitor-resistant cells to JQ1 and MGH-CP1, we performed ATAC-seq analysis. We examined changes in chromatin accessibility following treatment with BRD4 inhibitor, JQ1. The inhibitor treatment in A375 MEKi resistant cells induced extensive changes in chromatin state, as evidenced by the emergence of new regions with both significantly increased (’up-peaks’) and decreased (’down-peaks’) chromatin accessibility compared to the DMSO-treated control group **(Figure 5A)**. HOMER de novo motif analysis identified TEAD families as potential regulators associated with the down-regulated ATAC-seq peaks after JQ1 treatment. Additionally, Fra2, BATF, Fli1, STAT3, and IRF1 were identified as the main regulators associated with the up-regulated ATAC-seq peaks **(Figure 5B)**. These results suggest that BRD4 inhibition is associated with the downregulation of YAP1 effectors and the upregulation of immune regulators. Furthermore, GO analysis of specific accessible regions in the JQ1 treatment group revealed associations with response to dsRNA, immune response activating signaling, T cell co-stimulating signaling, leukocyte migration and other related processes. In contrast, specific accessible regions in the DMSO group were associated with positive regulation of biological process, response to stress, regulation of protein deacetylation and negative regulation of lymphocytes or T cell proliferation **(Figure 5C)**. These results emphasize the correlation between BRD4 inhibition and immune activation. Moreover, the chromatin accessibility of key target genes of YAP1, including *AXL*, *NRG1*, *BCL2L1* and *PD-L1*, decreased following treatment with either JQ1 or MGH-CP1 **(Figures 5D and S5A)**. Collectively, our results suggest that alterations in chromatin accessibility at YAP1 target genes and immune response genes are pivotal in mediating the effects of BRD4 or TEAD inhibition.

**Figure 5.**
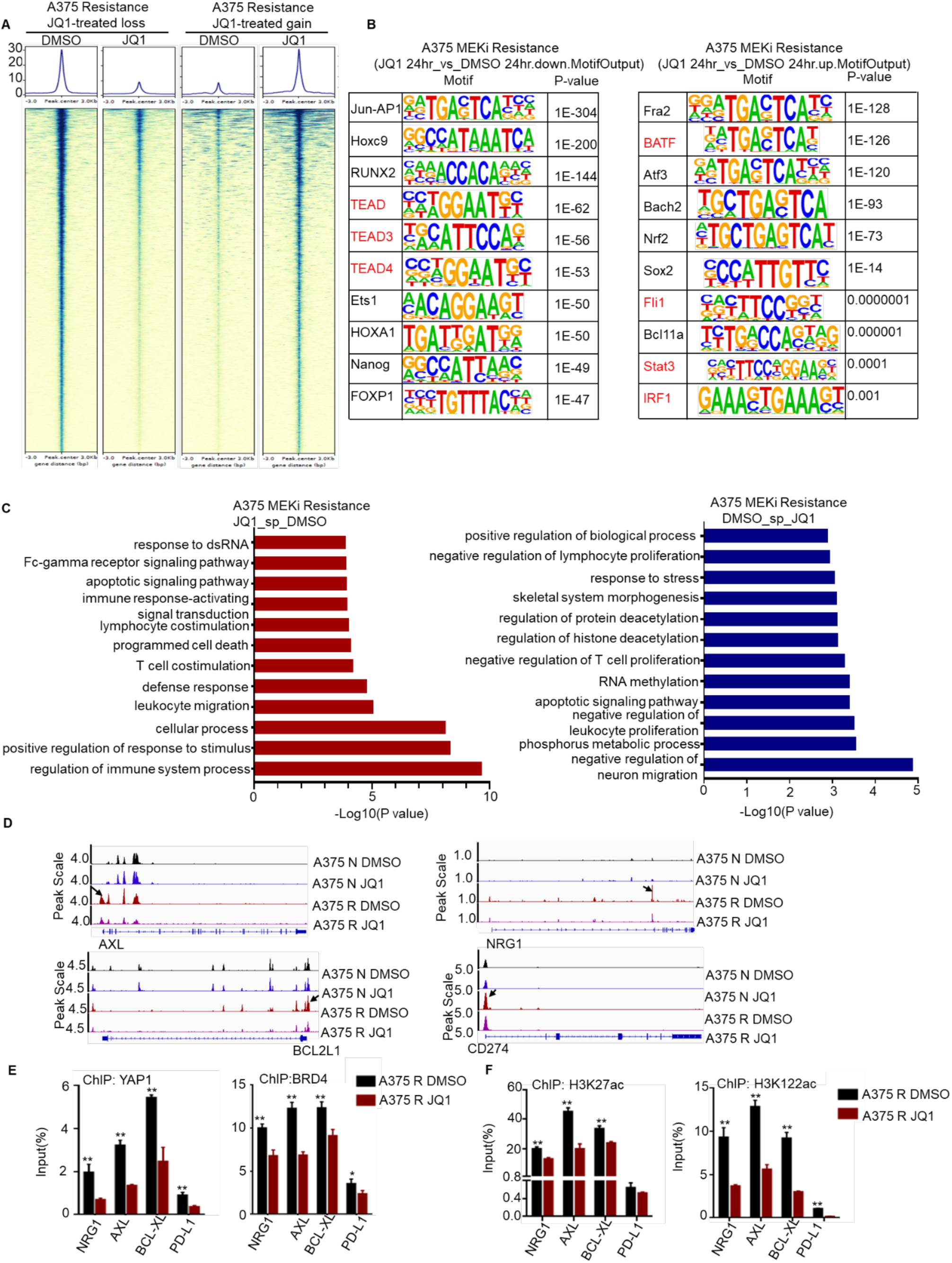
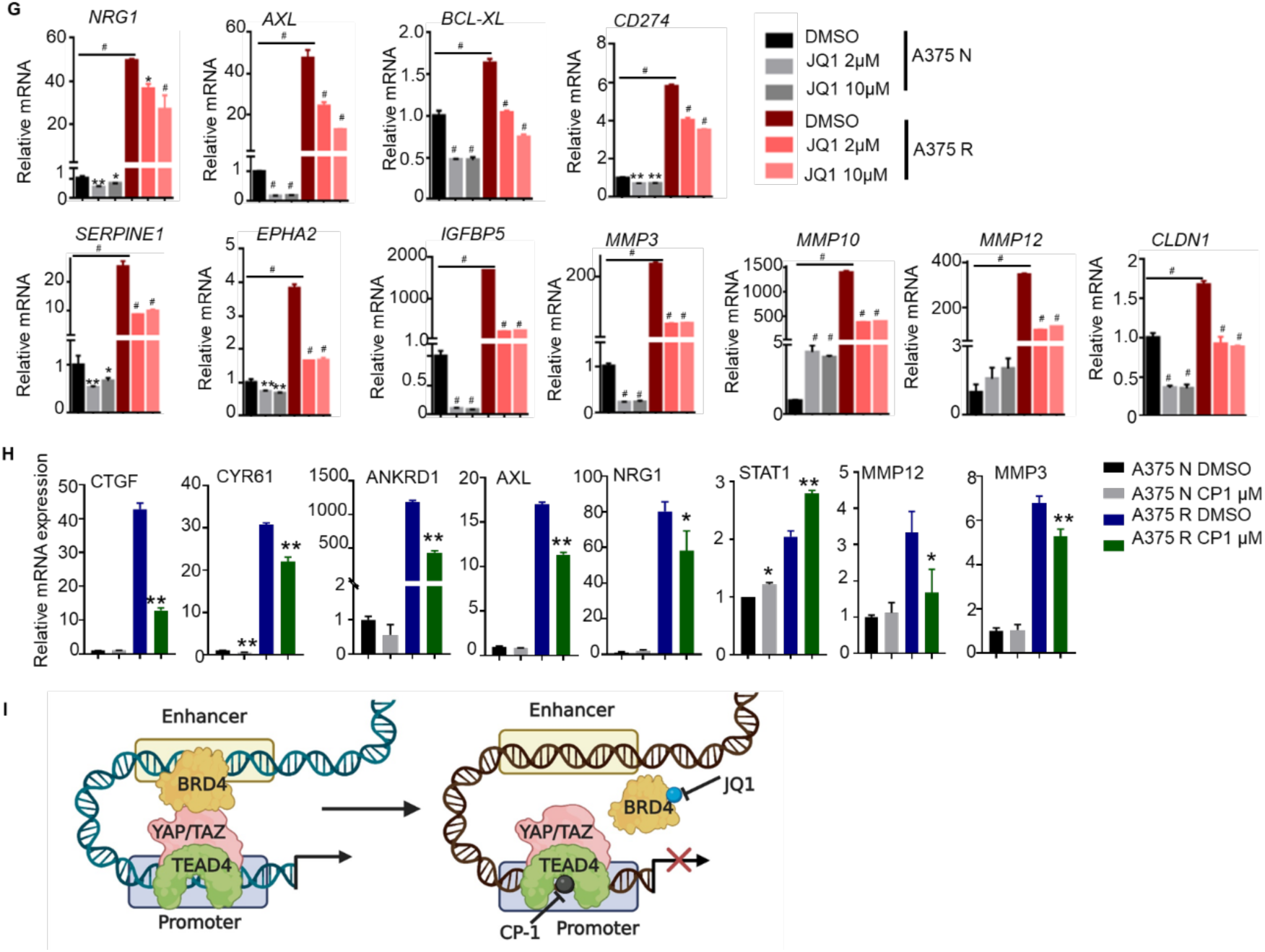
BRD4 or TEAD4 inhibition blunts the transcriptional expression of YAP1 target genes. A) ATAC-Sequencing analysis identifies gain or loss chromatin accessibility regions in MEKi-resistant cells after treatment with DMSO vs. JQ1. B) HOMER DNA-motif enrichment analyses of differentially accessible peaks in JQ1 vs. DMSO treatment group. The enrichment of binding motifs of 10 transcription factors per sample are shown. The p value is given. C) Gene ontology (GO) enrichment analyses showing distinct gene ontologies of target genes linked to differentially specific accessible peaks in JQ1 vs. DMSO. The x-axis indicates the −log10 (p-value). The y-axis indicates the names of enriched GO biological processes are indicated. D) Representative ATAC-sequencing tracks for the *AXL, BCL2L1, NRG1 and CD274* locus in A375 N and A375 R upon JQ1 treatment. The ATAC-seq data have been normalized to take sequencing depth into account and the scale on the y-axis was chosen for optimal visualization of peaks for each sample. E, F) ChIP-qPCR shows decreased YAP1, BRD4, H3K27ac, and H3K122ac binding on enhancers and promoters of YAP/TAZ targets (*NRG1*, *AXL*, *BCL-XL*, *CD274*) after JQ1 treatment in A375 MEKi resistant cells. ChIP with pre-immune IgG displayed background signal (which was comparable in all samples). DNA enrichment was calculated as % of input (n=3, *P<0.05, **P<0.01, ^#^P<0.0001). G, H) The relative mRNA expression level of YAP1 target genes was analyzed by qPCR after treatment with JQ1 or CP-1. The y axis shows the fold change in transcript levels versus DMSO-treated cells. Data are presented as mean ± S.D (n=3, *P<0.05, **P<0.01, ^#^P<0.0001). I) The schematic working model depicting the inhibition of BRD4 and TEAD in MEKi resistant melanoma cells (created with BioRender).

### 2.7 BRD4 or TEAD4 inhibition blunts the transcriptional level of YAP target genes

To investigate the impact of BRD4 inhibition on gene expression through the disruption of transcription factor binding, we conducted ChIP analysis. The results revealed a reduction in the binding of both YAP1 and BRD4 within the promoter loci of YAP1 target genes, including *NRG1*, *AXL*, *BCL*-*XL*, and *PD*-*L1* **(Figures 5E and S5B)**. Previous studies have highlighted that BRD4 is recruited not only to histones acetylated at H3K27, but also acetylates H3K122, leading to nucleosome eviction and chromatin de-compaction ^[21a]^. Similarly, we observed a striking decrease in the enrichment of H3K27ac and H3K122ac at the promoter regions of these genes following JQ1 treatment **(Figure 5F)**. These results suggest that YAP1 plays a crucial role in maintaining H3K122 acetylation in the target gene loci of BRD4, and the intrinsic acetyltransferase activity of BRD4 is necessary for YAP1’s transcriptional function. Furthermore, BRD4 inhibition led to reduced mRNA expression of YAP1 target genes, including *NRG1*, *AXL*, *BCL*-*XL*, and the immune suppressive gene *CD274* **(Figure 5G)**. Simultaneously, BRD4 inhibition decreased the expression of YAP1 target genes associated with proliferation (*SERPINE1*, *EPHA2*, and *IGFBP5*) and invasion (*MMP3*, *MMP10*, *MMP12*, and *CLDN1*) in MEKi resistant cells **(Figure 5G)**. Additionally, our data revealed that MGH-CP1 treatment of MEKi resistant cells, significantly decreased the expression of YAP1-TEAD target genes (*CTGF*, *CYR61*, *ANKRD1*, *AXL*, *NRG1, FST,* and *BCL2L1*), invasion genes (*MMP12*, *MMP3* and *MMP10*), and upregulated immune-active gene *STAT1* both MEK inhibitor resistant A375 and BRAF inhibitor resistant M14 melanoma cells **(Figures 5H and S5C)**. Analyzing published datasets ^[34]^, we found that TEAD inhibition, as anticipated, decreased the expression of YAP1 target genes, indicating the co-regulation of multiple target genes by TEAD and BRD4 **(Figure S5D)**. Together, these findings confirm that YAP1, TEAD, and BRD4 form a coordinated transcriptional complex that sustains oncogenic gene expression in resistant melanoma, and show that pharmacological inhibition with JQ1 or MGH-CP1 can effectively suppress this regulatory axis **(Figure 5I)**.

### 2.8 Expression levels of YAP/TAZ-TEAD are inversely related to the infiltration of cytotoxic T lymphocytes and the strength of the immune response

Emerging evidence suggests that the YAP1/TAZ-TEAD complex plays a role in promoting an immunosuppressive tumor microenvironment ^[36]^. In line with this, our data strongly suggests that elevated YAP1/TAZ-TEAD expression likely reprograms the expression of genes modulating an immune signature in vitro. To further define the coordination between YAP1 and TEAD in mediating immune evasion, we explored whether this correlation is mirrored in actual patient data. Our findings revealed that amplification of YAP1/TAZ copy number is associated with lower CD8+ T cell infiltration **(Figures 6A and S6A)**. Moreover, YAP1/TAZ expression is negatively correlated with cytotoxic T lymphocyte (CTL) numbers **(Figures 6B and S6B)**. Analyzing melanoma patient data from The Cancer Genome Atlas (TCGA), we found that YAP1/TAZ negatively correlates with antigen presentation and CD8+ T cell response genes while showing a positive correlation with the expression of immune suppressive genes such as *PD*-*L1*, *PD-L2*, *CD276*, *VTCN1*, and *CD47* **(Figure 6C)**. Notably, single-cell analysis of clinical melanoma tumors^[29b]^ indicates that, within the non-malignant compartment, YAP1, WWTR1, and TEAD4 are predominantly enriched in endothelial cells **(Figure 6D)**. Although these genes are generally recognized for their functions in tumor cells, their high expression in endothelial cells suggests that the tumor vasculature may actively contribute to immune evasion. To investigate the association between YAP1 activity and the immune landscape in melanoma, we analyzed transcriptomic data from patient-derived melanoma samples using XenoBrowser. Patients were stratified into YAP1-high and YAP1-low groups based on YAP1 expression levels. Heatmap analysis revealed that YAP1-high tumors exhibited an immune-suppressive, ‘cold’ tumor microenvironment in melanoma **(Figure 6E)**. This immune evasion phenotype was characterized by decreased antigen presentation (*HLA-A*, *HLA-B*, *TAP1*, *TAP2*, *TAPBP*), potentially impairing immune recognition, and a suppressed CD8⁺ T cell response, as evidenced by reduced expression of cytotoxic markers (*GZMA*, *GZMB*, *PRF1*, and *IFNG*). Moreover, YAP1/TAZ-high tumors exhibited increased expression of immune checkpoint molecules (*PD-L1*, *PD-L2*, *CTLA4*, *TIM3*, *LAG3*), which contribute to T cell exhaustion. The cytokine and chemokine profile further reflected an immunosuppressive state, with decreased levels of pro-inflammatory signals (*CXCL9*, *CXCL10*, *IFI27*) and increased expression of the M2 macrophage marker *CD206*, suggesting a tumor-promoting macrophage polarization **(Figure 6E)**. A similar pattern was observed for WWTR1 (TAZ) expression, which showed comparable correlations with immunosuppressive gene signatures **(Figure S6C)**. Collectively, these results indicate that elevated activity of YAP1 and WWTR1 is associated with an immunosuppressive tumor microenvironment, potentially contributing to immune evasion and resistance to immunotherapy in melanoma. Notably, *PD*-*L1*, an immune inhibitory checkpoint, exhibited significantly higher expression levels in metastatic melanoma compared to primary melanoma **(Figure 6F)**. Importantly, YAP1 and WWTR1 showed a significant positive correlation with *PD-L1* in metastatic melanoma **(Figures 6G and S6D)**, and knock down of YAP1 in MEKi resistant cells resulted in increased expression of immune responsive genes *IL*-*24*, *STAT1*, and *CXCL10* **(Figure 6H)**. These observations indicate that YAP1/TAZ-TEAD is negatively associated with components of an active immune response, thereby promoting immune escape and tumor progression.

**Figure 6.**
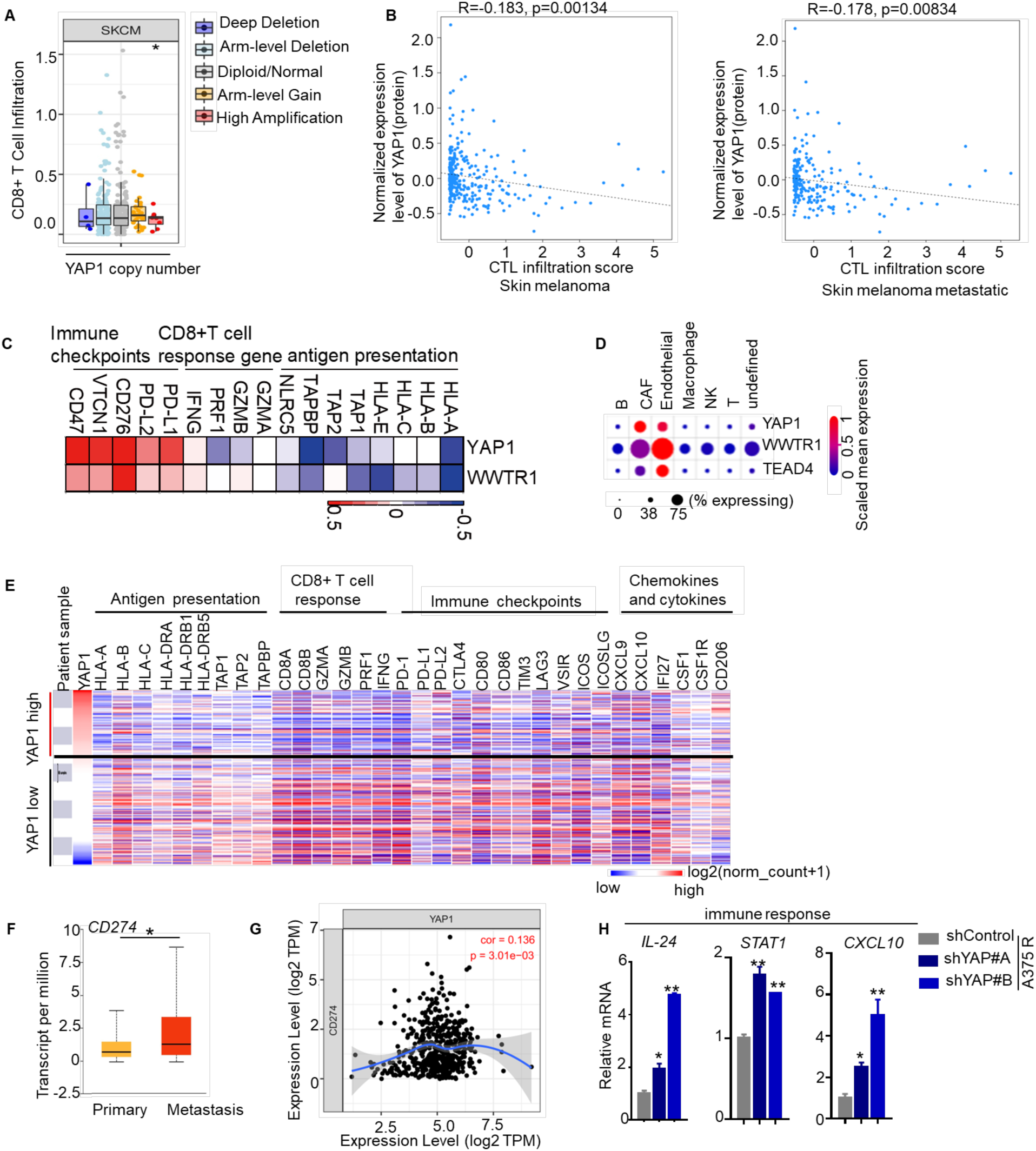
YAP1 expression is negatively correlated with cytotoxic T lymphocytes infiltration and immune response (TCGA dataset). A) Box plots are presented to show the distributions of CD8+ T cell with different somatic copy number aberrations of YAP1. B) Scatterplot displaying a negative correlation between YAP1 and cytotoxic T (CTL) infiltration. C) The heatmap depicts the Spearman’s correlation coefficients between YAP1 and WWTR1 expression and genes associated with antigen presentation, CD8⁺ T cell function, and immune suppression. Gene expression data were obtained from The Cancer Genome Atlas (TCGA). D) Heatmap displays the expression levels of YAP1, WWTR1, and TEAD4 across different cell types, including B cells, CAFs (cancer-associated fibroblasts), endothelial cells, macrophages, NK cells, T cells, and undefined cells. The dot size represents the scaled mean expression of each gene, with the color gradient from blue (low expression) to red (high expression). The data was based on the analysis from Jerby-Arnon, L. et al. E) Xena Browser Visual Spreadsheet displays YAP1, antigen presentation genes, CD8+ T cell signature genes, immune checkpoint, cytokines, and chemokines in melanoma patients. The expression is colored red to blue for high to low expression. F) The bar graph indicates the expression level of *CD274* in primary (n=104) and metastasis (n=368) SKMC (*P<0.05). G) The scatter plot shows the correlation between *YAP1* and *CD274* in SKMC tumor (103 samples). H) The relative mRNA expression level of immune genes was analyzed by qPCR after YAP1 knock down in MEKi resistant cells. The y axis shows the fold change in transcript levels versus DMSO-treated cells. Data are presented as mean ± S.D (n=3, *P<0.05, **P<0.01).

### 2.9 TEAD inhibitor exerts an anti-tumor effect and significant survival benefit in xenograft mouse model by reactivating immune response genes

Since YAP1-TEAD transcriptional activity regulates the expression of genes critical for immune suppression, we conducted an integrated analysis using TCGA, TIMER, and additional cohort datasets from the TIDE analysis platform to evaluate the role of the TEAD family in melanoma. Our findings revealed a significant association between elevated expression of TEAD1, TEAD2, TEAD3, and TEAD4 and poorer overall survival in melanoma patients **(Figures 7A and S7A)**. Furthermore, TEAD3 and TEAD4 expression exhibited a strong inverse correlation with cytotoxic T-cell infiltration in metastatic melanoma, as observed in both the GSE65904 dataset and TCGA **(Figure 7B)**. Similarly, TEAD1 and TEAD2 expression also negatively correlated with cytotoxic T-cell infiltration in metastatic melanoma from TCGA **(Figure S7B)**. Notably, the gain or amplification of TEAD gene copy numbers (TEAD1, TEAD2, TEAD3, TEAD4) is associated with lower CD8⁺ T cell infiltration in melanoma, implying that TEAD copy number alterations may contribute to immune evasion and potentially facilitate melanoma progression by impairing immune surveillance **(Figure S7C)**. Moreover, pharmacological inhibition of TEAD activity using MGH-CP1 significantly reduced *PD-L1* expression in MEK inhibitor-resistant cells compared to their parental counterparts **(Figures 7C and S7D)**, suggesting a potential role for TEAD signaling in immune evasion. Given that TEAD inhibitors are currently being evaluated in clinical trials for advanced mesothelioma and NF2-mutant cancers^[37]^, we sought to determine whether TEAD inhibition could suppress tumor growth and enhance anti-tumor immune responses in melanoma in vivo. To this end, we evaluated the therapeutic efficacy of combining CP-1 with anti-PD-1 immune checkpoint blockade, we treated D4M-UV3^[38]^ melanoma-bearing mice with vehicle, CP-1 alone, anti-PD-1 alone, or the combination of CP-1 and anti-PD-1. Kaplan-Meier survival analysis revealed that both CP-1 and anti-PD-1 monotherapies moderately prolonged survival compared to the vehicle group **(Figure 7D)**. Notably, the combination treatment resulted in a marked improvement in survival, with a substantial proportion of mice remaining tumor-free throughout the observation period **(Figure 7D)**. Statistical comparison confirmed that the combination therapy significantly outperformed either monotherapy, indicating a strong synergistic effect between CP-1 and immune checkpoint inhibition **(Figure 7D)**, while having no detectable impact on mouse body weight, indicating treatment tolerability **(Figure S7E)**. Infiltration of CD8⁺ T cells was significantly increased in both the MGH-CP1, PD-1 and combination treatment groups, **(Figures 7E)**, suggesting that PD-1 blockade helps mitigate the immune-suppressive environment in melanoma, and that MGH-CP1 treatment further enhances this effect. These findings highlight the crucial role of TEAD in mediating drug resistance and immune evasion in melanoma, with significant clinical implications. In conclusion, inhibiting TEAD can disrupt YAP-dependent tumor initiation and growth in vivo, offering potential therapeutic benefits in overcoming immune evasion and resistance mechanisms.

**Figure 7.**
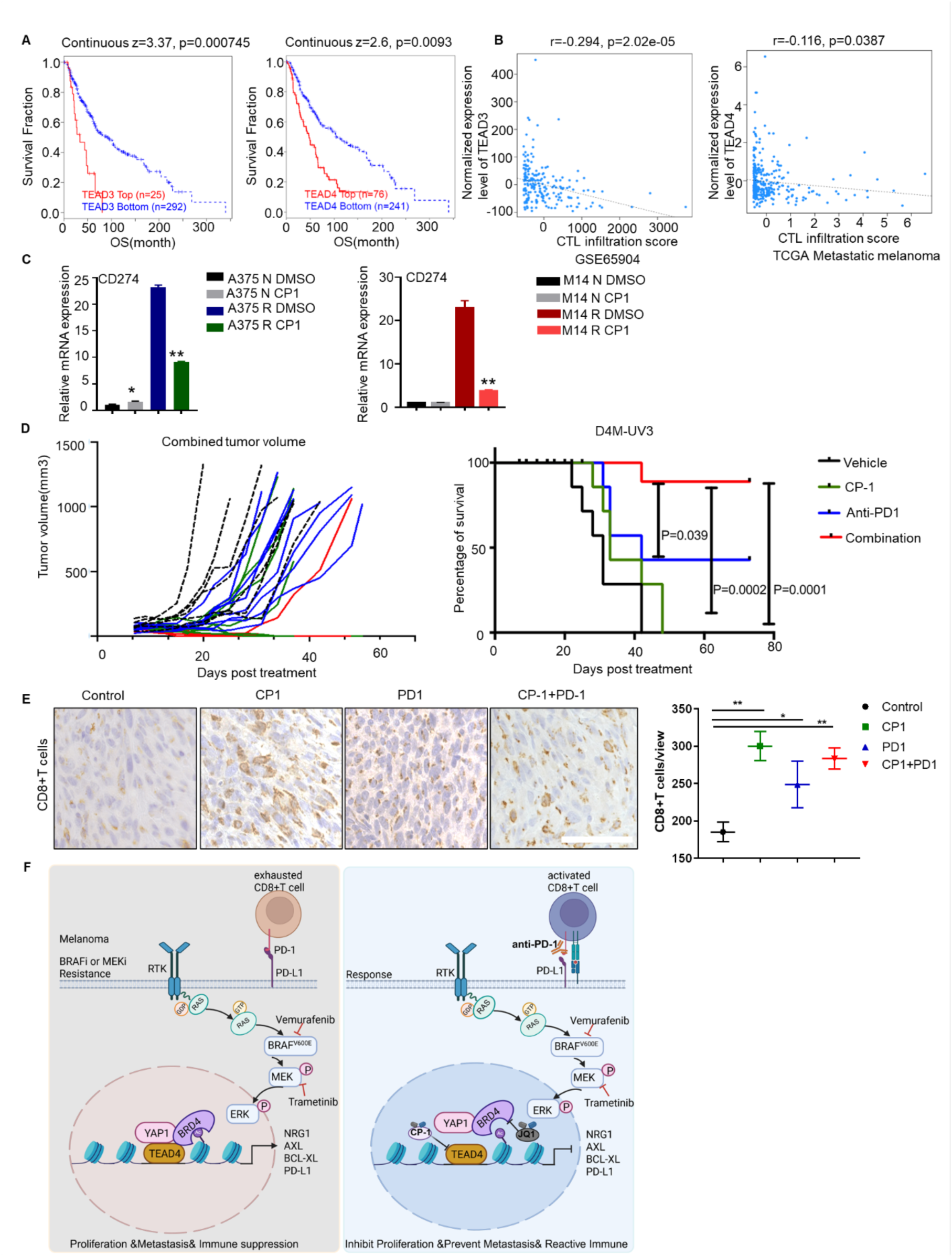
TEAD inhibitor exerts an anti-tumor effect and significant survival benefit in xenograft mice model by reactivating immune response. A) The OS of TEAD3 and TEAD4 in SKMC patients from TCGA were analyzed by Kaplan-Meier plotter. B) The scatterplot displaying the correlation between TEAD3/TEAD4 and cytotoxic T (CTL) infiltration score. C) mRNA-expression level of PD-L1 after 24 h treatment by JQ1 or MGH-CP1. Values are normalized to housekeeping gene 18s. Data represents as mean ± s.d. after normalization to DMSO-treated cells. *P<0.05; **P<0.01. D) The tumor growth curve and Kaplan Meier survival curve of melanoma in mice treated with Vehicle (n=7), CP-1 (n=8), anti-PD-1 (n=7), and CP-1 plus anti-PD-1 (n= 7). Right: Immunohistochemistry (IHC) staining of CD8⁺ T cells and YAP1 in tumors isolated from mice with the corresponding treatment. Scale bar: 100 µm. Left: Statistical analysis of CD8⁺ T cell numbers in each field of view under the macroscope (n=4). F) Schematic of crosstalk among YAP1/TEAD4/BRD4 complex in BRAF mutant melanoma. In BRAF mutant melanoma, the BRAF signal pathway and YAP1/TEAD4/ BRD4 cooperate to regulate melanoma growth, the YAP1/TEAD4/BRD4 complex exhibits higher transcriptional activity in BRAF inhibitor resistant melanoma cells. Targeting BRD4 or TEAD4 by JQ1 or CP-1 blunts YAP1/TEAD4/BRD4 complex transcriptional activity and inhibits BRAF inhibitor melanoma growth (created with BioRender).

In summary, our findings strongly suggest that the YAP1/BRD4/TEAD axis plays a critical role in MEK inhibitor-resistant melanoma. We show that knockdown of YAP1 enhances the expression of anti-tumor cytokines and chemokines while decreasing immune-suppressive gene expression. Moreover, YAP1 amplification is linked to reduced cytotoxic T-lymphocyte (CTL) infiltration and a ‘cold’ immune response. Importantly, inhibiting BRD4 or TEAD4 blocks YAP transcriptional activity and reactivates a ‘hot’ immune profile. Collectively, our results highlight the significant crosstalk between the YAP1-TEAD/BRD4 axis and MAPK signaling, suggesting a promising combination strategy to improve the therapeutic efficacy of RAF or MEK inhibitors, as well as immune checkpoint inhibitors (anti-PD1), in melanoma patients **(Figure. 7F)**.

## 3. Discussion

The high prevalence of BRAF mutations, particularly the V600E variant, in melanoma underscores the critical importance of targeting the MAPK signaling cascade therapeutically^[39]^. However, despite initial clinical successes with RAF and MEK inhibitors (BRAFi/MEKi), both intrinsic and acquired resistance inevitably emerge, significantly limiting long-term treatment efficacy^[40]^. Similar challenges in overcoming MAPKi resistance have been extensively documented across multiple BRAF-mutant malignancies, including non-small cell lung cancer, colorectal, and thyroid cancers^[41]^. While prior studies have linked activation of the Hippo-YAP1 pathway to MAPKi resistance ^[2c, 4]^, our study substantially expands this concept by elucidating the molecular mechanisms through which YAP1 coordinates chromatin-centric transcriptional reprogramming and immune evasion. Unlike prior reports that focused on YAP1 upregulation or nuclear translocation as markers of resistance^[42]^, our findings highlight the functional dependence on downstream YAP/TEAD/BRD4 mediated transcription, independent of YAP1 overexpression. This distinction is critical, as it suggests that resistant melanoma cells sustain their adaptive transcriptional state through chromatin reprogramming and transcriptional activation, rather than through upstream signaling alone.

While YAP fusion-driven condensates recruit BRD4/TEAD in ependymoma ^[43]^, our findings show that endogenous YAP1 similarly engages TEAD and BRD4 to sustain resistance in melanoma. Moreover, although TEADs have been linked to invasive melanoma states ^[44]^, we demonstrate that (BRAFi/MEKi)-resistant cells are uniquely sensitive to TEAD and BRD4 inhibition, revealing a distinct therapeutic vulnerability. These studies support our findings and highlight the translational relevance of targeting YAP/TEAD/BRD4 complexes in resistant melanoma. Our integrated clinical and preclinical analyses robustly identify YAP1 activation as a defining hallmark of BRAFi/MEKi resistance. Elevated expression of YAP1 target genes in biopsies from resistant melanoma patients strongly correlates with aggressive tumor characteristics. Mechanistically, YAP1 directly engages with BRD4 and TEAD transcription factors, orchestrating extensive chromatin remodeling to sustain oncogenic transcriptional activity while simultaneously repressing critical genes required for effective immune responses. Thus, YAP1 emerges as a pivotal nexus linking treatment resistance and immune suppression. Supporting this mechanistic link, we show that pharmacological disruption of BRD4 or TEAD function effectively reverses YAP1-driven transcriptional signatures, facilitating the restoration of cytotoxic T-cell infiltration into the tumor microenvironment. Notably, the preferential sensitivity of resistant tumor cells to BRD4 and TEAD inhibitors further implies that dependency on the YAP1-BRD4-TEAD complex evolves adaptively under prolonged MAPKi treatment, representing a potential therapeutic vulnerability.

Highlighting the clinical relevance of targeting the YAP1-TEAD axis, our study demonstrates the efficacy of the palmitoylation-dependent TEAD inhibitor, MGH-CP1. Unlike earlier nonspecific inhibitors of the YAP-TEAD interaction, such as verteporfin^[45]^, MGH-CP1 selectively disrupts TEAD auto-palmitoylation, effectively destabilizing YAP-TEAD complexes and suppressing downstream oncogenic transcriptional outputs ^[35a]^. Importantly, in mice models, MGH-CP1 treatment not only inhibits tumor progression but also reinvigorates antitumor immunity through enhanced infiltration and activation of CD8^+^ T-cells. These immune-stimulatory effects were notably amplified by combination with anti-PD-1 checkpoint blockade therapy. Our findings are consistent with recent study showing that YAP1 is a key mediator of anti-PD-1 resistance, where pharmacological YAP1 inhibition with verteporfin restored anti-PD-1 responsiveness and promoted a pro-inflammatory tumor microenvironment in YUMM1.7 melanoma models^[46]^. Furthermore, analysis of melanoma patient sample revealed a significant inverse correlation between YAP/TAZ-TEAD signaling activity and markers of cytotoxic immune infiltration, strongly suggesting that TEAD inhibition could effectively transform immunologically “cold” resistant tumors into immune-reactive “hot” microenvironments.

A critical mechanistic question arising from our study relates to how YAP1 dynamically coordinates BRD4 and TEAD recruitment to chromatin regions during therapy resistance. BRD4 binds preferentially to acetylated histone marks, particularly H3K27ac, which we observed enriched at YAP1-targeted loci in resistant cells. This observation supports a putative feedforward loop wherein YAP1-TEAD complexes recruit BRD4 to sustain acetylation-dependent chromatin accessibility, facilitating persistent oncogenic transcription. This hypothesis is further reinforced by the efficacy of BET inhibitor JQ1 in suppressing YAP1-driven gene expression. Future studies mapping the precise interplay among YAP1, BRD4, TEAD, and chromatin remodeling complexes will be instrumental in refining combination therapeutic strategies.

In summary, our study redefines YAP1 as a central chromatin-immune regulator that bridges MAPKi resistance and immune evasion through its cooperative interactions with BRD4 and TEAD. Targeting the YAP1-BRD4-TEAD axis presents an innovative paradigm shift from sequential single-agent therapies toward integrative epigenetic-immunological combination strategies. Ultimately, this approach holds considerable promise for improving clinical outcomes in patients with aggressive, immune-evasive melanoma.

## 4. Experimental Section

### Data and code availability

The analysis of Hippo pathway related genes was obtained from TCGA, the analysis of the expression level of YAP1 and BRD4 in SKCM primary and metastasis tumors was based on previous published data^[47]^. The analysis of YAP1 target gene expression in patient sample was obtained from UALCAN^[48]^. The overall survival analysis of melanoma patients with YAP1 or BRD4 expression level was from Tumor Immune Dysfunction and Exclusion (TIDE) website^[26]^. ATAC-seq and RNA-seq data have been deposited to in the NCBI Gene Expression Omnibus database and is publicly available as of the date of publication. This paper does not report original code. Any additional information required to reanalyze the data reported in this paper is available from the lead contact Srinivas Vinod Saladi.

### Cell lines, chemicals, and treatments

All cell lines were cultured at 37 °C in the presence of 5% CO2. HEK293T, A375 parental and A375-Dabrafenib resistant cells were cultured in DMEM (Lonza#: BE12-604F); M14 parental and M14-PLX4720 resistant cells were maintained in RPMI1640 (Catalog#: 10-040-CV, Corning). All the mediums were supplemented with 10% fetal bovine serum (FBS) and 100 units/ml antibiotic antimycotic solution (Catalog #: A5955, Sigma-Aldrich). JQ1 (Catalog#: SML1524) and OTX-015 (Catalog#: SML1605) were purchased from Sigma-Aldrich. MGH-CP1 was a gift from Dr. Xu Wu. All inhibitors were dissolved in DMSO. Cells were treated with JQ1, OTX015, GSK-126, MS-275, verteporfin and MGH-CP1 by varied concentrations and times as indicated.

### In-vivo experiment

The C57BL/6 syngeneic mouse melanoma cell line D4M.3A.3-UV3 cells were cultured in DMEM with 1% penicillin/streptomycin/l-glutamine and 10% FBS as previously described ^[49]^. For syngeneic mouse models, 6 to 8-week-old female C57BL/6 mice were obtained from Jackson Laboratory. One million melanoma D4M.3A.3-UV3 cells in PBS were inoculated subcutaneously in the right flank. Vehicle (5% v/v PEG-400 and 5% v/v Tween-80 in PBS) or MGH-CP1 daily i.p. at 75 mg/kg was administered intraperitoneally daily for the duration of the experiment, starting 6 days after tumor reaching to 50-100 mm^3^. Blocking antibodies, anti–PD-1 (a gift from Gordon Freeman, Dana-Farber Cancer Institute, Boston, MA) was administered intraperitoneally on days 7, 9, and 11 at a dose of 200 μg per mouse. For survival studies, mice were sacrificed when tumors reached a maximum volume of 1,000 mm^3^. Tumor volumes were measured using digital calipers and calculated by the following formula: volume (mm^3^) = (width^2^ × length)/2.

### Cell Viability Assays

Cells (1.5X10^3^/well) were seeded into 96-well plates and treated the next day with different concentrations of indicated drugs. After 72h treatment, cell viability was measured by CellTiter-Glo® Luminescent assay (Catalog #G7570, Promega) according to the manufacturer’s protocol. Luminescence in the plate was measured using a microplate reader (SpectraMax M5, Molecular Devices, Sunnyvale, CA, USA). Percent cell growth was calculated relative to DMSO treated cells. Dose-response curve was then plotted as percentage viability against the log concentration of indicated drugs. IC50 was determined using a sigmoidal regression model using GraphPad Prism 7.0.

### Lentiviral Infection

HEK293T cells were used to package virus. 2 × 10^7^ cells were plated in 10 cm tissue culture dish and transfected with 10μg of the pLKO.1 or pLKO.1 lentiviral vector encoding shYAP1, 10 μg of pVSV-G, 10 μg of pRGR and 10 μg of pRSV with Calcium phosphate transfection kit (Catalog #:631312, Takara) according to the manufacturer’s instruction. Conditioned medium containing recombinant lentiviruses was collected at 36 hours after transfection and filtered through non-pyrogenic filters with a pore size of 0.45 μm (Catalog#: SLHP033RS, Millipore, Sigma). Samples of these supernatants were applied immediately to target cells together with Polybrene (Catalog #:H9268, Sigma-Aldrich) at a final concentration of 10 μg/ml, and supernatants were incubated with cells for 24 h. After infection, cells were placed in fresh growth medium and cultured as usual. The infected cells were harvest after 72-80 h after infection for further analysis.

### RNA Extraction and Quantitative PCR (QPCR)

Total RNA of the cells treated by JQ1 and shYAP1 was extracted using TRIzol: Chloroform phase-separation by centrifugation followed by RNA precipitation using isopropanol, or RNeasy Plus Mini Kit (QIAGEN, Catalog# 74134) according to the manufacturer’s instructions. The RNA concentrations were measured with Nanodrop (Thermo Fisher Scientific) and 1 μg of total RNA was used for cDNA synthesis using the Applied Biosystems™ High-Capacity cDNA Reverse Transcription Kit with RNase Inhibitor (Catalog# 4374967, Thermo Fisher Scientific). The qPCR reactions were set up in 20 μL using Power SYBR™ Green PCR Master Mix (Thermo Fisher Scientific, Catalog#4368702) according to manufacturer’s instructions and 2 μL of 1:10 diluted cDNA. The reactions were run in Quant Studio 6 Flex Real-Time PCR Systems (Applied Biosystems). The gene expression levels were normalized to 18s housekeeping gene expression levels in each sample. The data were presented as fold changes of gene expression in the JQ1 treated sample compared to the control.

### RNA-seq

A375 parental and A375 MEK inhibitor resistant cells were incubated in biological triplicates for 72 hr with shControl, shYAP1#A, and shYAP1#B treatment. Total RNA was extracted using RNeasy Plus Mini Kit (QIAGEN, Catalog# 74134). RNA concentrations were measured and quality controlled on a Bioanalyzer, RNA-Seq libraries were made and sequenced using NovaSeq PE150.

### Immunoblotting

A375 and M14 parental (N) and BRAFi/MEKi-resistant cells were lysed in RIPA buffer supplemented with protease and phosphatase inhibitors (Roche). Protein concentrations were determined using the DC™ Protein Assay Kit (Bio-Rad), denatured in 2× SDS loading buffer at 95 °C for 5 min, and resolved by SDS-PAGE (4–20% gradient gels, Bio-Rad). Proteins were transferred to PVDF membranes (Millipore), blocked in 5% blocking buffer (Bio-Rad) for 1 h, and incubated overnight at 4 °C with primary antibody PD-L1 (Cat. GTX104763; GeneTex). After washing, membranes were incubated with HRP-conjugated secondary antibodies (CST) and imaged using the iBright CL750 system (Thermo Fisher Scientific).

### ATAC-seq

The ATAC (Assay for Transposase-Accessible Chromatin) method was applied based on the published protocol ^[50]^. A375 parental and A375 Dabrafenib resistant cells were plated at 5 x 10^6 cells into 10 cm plates, and treated the next day with DMSO or 2 µM or 10 µM JQ1 or MGH-CP1.

Then the cells were trypsinized at 24 h. After all samples were harvested, 50000 cells /sample were resuspended in 1 mL of cold PBS. Cells were centrifuged at 500g for 5 min in a pre-chilled (4°C) fixed-angle centrifuge. After centrifugation, the supernatant was carefully completely aspirated. Cell pellets were then resuspended in 50 μL of ATAC-seq RSB containing 10 mM Tris-HCl pH 7.4, 10 mM NaCl, and 3 mM MgCl2 0.1% NP40, 0.1% Tween-20, and 0.01% digitonin by pipetting up and down three times. This cell lysis reaction was incubated on ice for 3 min. After lysis, 1 mL of ATAC-seq RSB containing 0.1% Tween-20 (without NP40 or digitonin) was added, and the tubes were inverted to mix. Nuclei were then centrifuged for 5 min at 500g in a pre-chilled (4°C) fixed-angle centrifuge. The supernatant was removed and nuclei were resuspended in 50 μL of transposition reaction: 2.5 μL Tagment DNA Enzyme 1 (TDE1) (Illumina, Catalog#15027865), Tagment DNA Buffer (Illumina, Catalog #15027866), 16.5 μL PBS, 0.5 μL 1% digitonin, 0.5 μL 10% Tween-20, and 5 μL water by pipetting up and down five times. Transposition reactions were incubated at 37°C for 30 min in a thermomixer with shaking at 1,000 r.p.m. Reactions were cleaned up with DNA Clean & Concentrator-5 Kit (Zymo research, Catalog#D4013). Quality control of tagmented DNA was performed by TapeStation. Libraries were amplified as described previously ^[50]^. 50-bp paired-end reads were generated on a Nextseq instrument (Illumina).

### ATAC-seq analysis

Paired-end fastq files were trimmed of adapter sequence using NGMerge ^[51]^ and mapped to the GRCh38 genome assembly using Bowtie2.3.4.3 ^[52]^ using the following flags: –-very-sensitive and –k20. For normalized bigwig pileup diagrams, aligned reads were normalized to CPM using the bamCoverage tool in deepTools ^[53]^ with a binSize of 10. Differential peaks were identified using csaw v1.7.1^[54]^, first by filtering for a log2 fold change of >2 using windows of 2000bp, and computing differential peaks using edgeR v3.32.1^[55]^. Peaks were annotated by known genes within 3000 bp upstream of the TSS and 1000 bp downstream of the TES.

### ChIP-assay

For inhibitor treatment, A375 parental and A375-Dabrafenib resistant cells were plated at 5 x 10^6^ cells into 10 cm plates, and were treated the next day either with DMSO or with 10 µM JQ1. For YAP1 knock down, A375 parental and A375-Dabrafenib resistant cells were plated at 5 x 10^5^ cells into 10 cm plates, and were infected with shControl or shYAP1. The procedure was followed as previously described ^[56]^. Briefly, cells were trypsinized after 24 hours of treatment or 80 hours of infection. Cells were washed in phosphate-buffered saline (PBS) and crosslinked with 1% Formaldehyde for 10 min or crosslinked with two agents starting with 2 mM EGS (ethylene glycol bis (succinimidylsuccinate)) for 30 min at RT, followed by 1% formaldehyde for 10 min. Crosslinked cell lines were quenched with 125mM glycine for 5 minutes at room temperature. After quenching, the fixed cell pellets were suspended in RIPA buffer and rotated for 3-4 hours at 4°C, nuclear lysates were sonicated 8 minutes for 5 times in a bioruptor (Diagenode). Soluble chromatin was immunoprecipitated with 4 μg of H3K27ac, H3K122ac or IgG antibody; or 2 μg of YAP1, BRD4, WWTR1, TEAD4 or IgG antibody. Precipitated DNA was purified by DNA Clean & Concentrator-5 Kit (Zymo research, Catalog#D4013). Quantitative PCR was performed as previously described.

### Ethics statement

All studies and procedures involving animal subjects were approved by the Institutional Animal Care and Use Committees of Massachusetts General Hospital (Boston, MA) and were conducted strictly in accordance with the approved animal handling protocols. The maximal tumor volume of animal experiments was one-tenth of body weight and all experiments did not exceed this limit.

### Statistical analysis

Statistical analyses were performed utilizing GraphPad Prism 7.0. All statistical details of the experiments, including exact values of n and what n represents, are mentioned in the figure legends. For quantitative reverse transcription PCR data expressed as relative fold changes, Student’s *t* test and one-way analysis of variance (ANOVA) with Dunnett’s test were used for pairwise comparisons and multigroup comparison, respectively. Data are presented as means ± standard deviation (SD). ^∗^p values ≤0.05 were considered statistically significant, with p ≤ 0.01 designated ^∗∗^, p ≤ 0.001 designated ^#^.

## Acknowledgments

We thank the Mass General Hospital next-gen sequencing core for the help with the sequencing. We are grateful to members of the Saladi laboratory for comments on the manuscript and technical help with the experiments. DEF expresses his gratitude to the Lancer Family for their support of the Lancer Endowed Chair at Harvard Medical School. This work was supported by Melanoma Research Foundation awarded to GB and SVS, Normal Knight Leadership development funds and Mike Toth Head and Neck Cancer Center funds to S.V.S. DEF gratefully acknowledges research support from the Dr. Miriam and Sheldon Adelson Medical Research Foundation, NIH (2R01AR043369-27, 5P01CA163222, 2R01AR072304-06), and the US Department of Defense (Melanoma Academy grant W81XWH2220052 and grant HT94252310948).

## Data Availability Statement

Sequencing data have been deposited with Gene Expression Omnibus (GEO) and are publicly available as of the date of publication. Accession numbers are listed in the key resources table. This paper does not report original code. Any additional information required to reanalyze the data reported in this paper is available from the lead contact upon request.

## Author contributions

S.V.S. conceptualized, designed, and supervised the study. N.C. designed the study, performed experiments, and interpreted data. R.I.S., D.E.F., and S.V.S. supervised the study. N.C., G.B., B.L., and R.I.S., performed computational analysis and interpreted data. H.B., M.S., G.B., I.D.S., X.W., D.E.F. designed the study and interpreted data. H.B., N.A., M.S., A.M.E., K.S.E., and R.U. analyzed and interpreted data. A.A.E., Y.S., designed and performed experiments. Y.S. synthesized MGH-CPI. M.U provided technical assistance under the supervision of N.C., and S.V.S. N.C., and S.V.S. wrote the manuscript. All authors approved the final submitted manuscript.

