## Supplementary data for "YAP1 drives aggressive and therapy resistant state in melanoma through reprogramming the chromatin and regulating immune evasive programs"

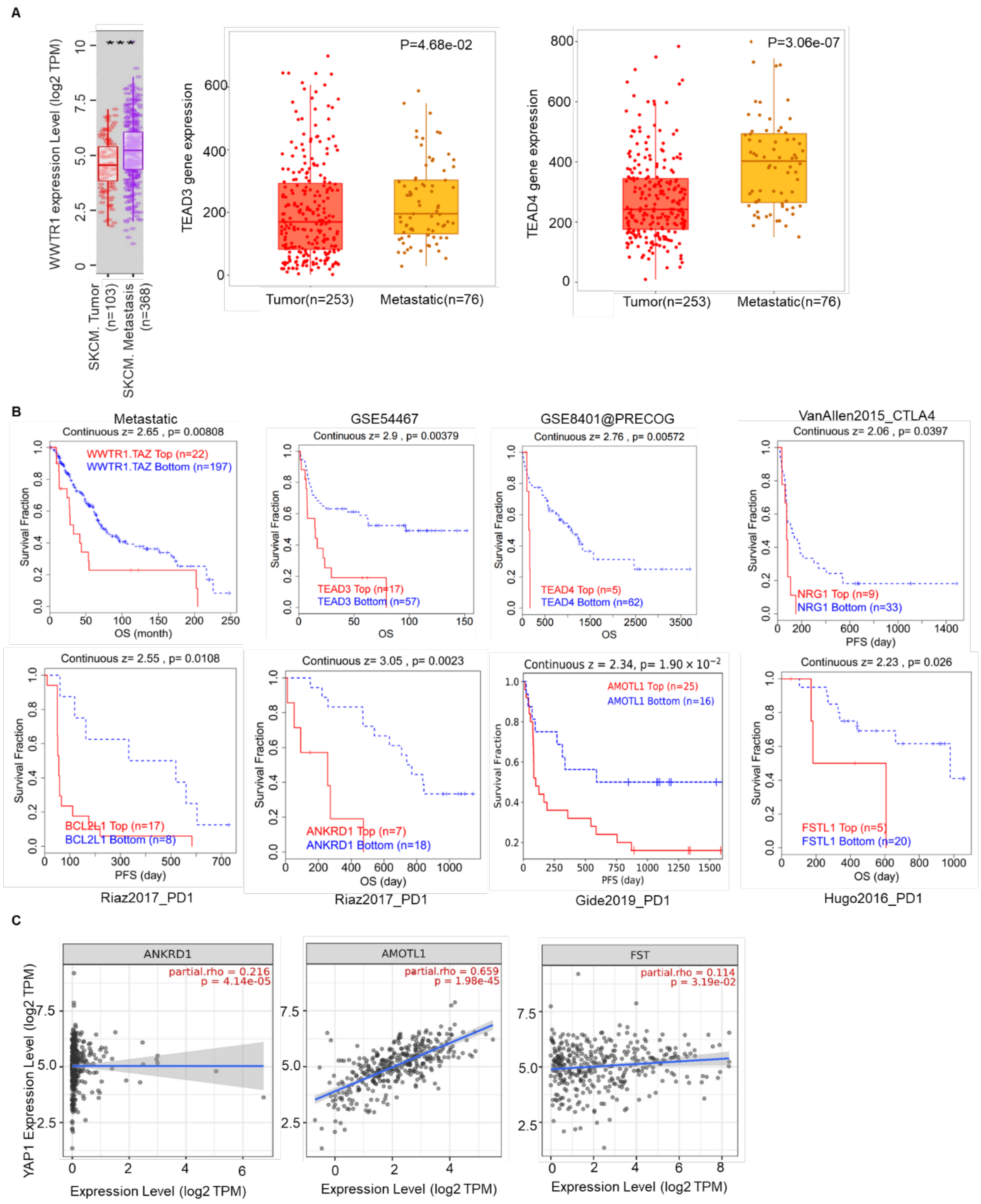


**Figure S1. Hippo pathway components are highly expressed in metastatic melanoma and correlated with poor survival.** A) Analysis of differential gene expression (DiffExp) of *WWTR1*, *TEAD3* and *TEAD4* in SKCM primary and metastasis patients. ****P*<0.001. B) The overall survival (OS) analysis of *WWTR1(TAZ)*, *TEAD3* and *TEAD4* and their target genes (*NRG1*, *BCL2L1*, *ANKRD1*, *AMOTL1*, *FSTL1*) in melanoma. C) The correlation of between *YAP1* and *ANKRD1*, *AMOTL1*, and *FST* in metastatic melanoma.

A

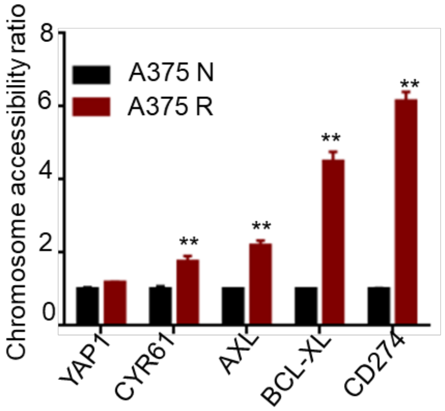


**Figure S2. ATAC-analysis identified differentially chromatin accessibility landscape in melanoma cells.** A) qPCR analysis of genomic DNA fragments released from Tn5 transposase-digested nuclei lysed from A375 N and A375 R. qPCR was carried out using primers that target open chromatin regions, qPCR data were normalized to the GAPDH promoter site. Data are shown as mean±SD (n=3, **P<0.01).

**A**

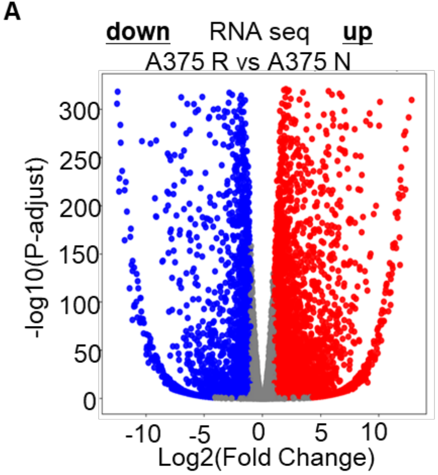


**Figure S3. Activation of YAP1 Transcriptional program is involved in MEK inhibitor resistance.** A) Differential expression of RNA-seq analysis in A375 Native and A375 Resistant (with acquired resistance to MEK inhibitor) cells. Volcano plot indicates the significance of differentially expressed genes (DEGs). Red indicates the genes with up-regulation; Blue indicates down-regulated genes.

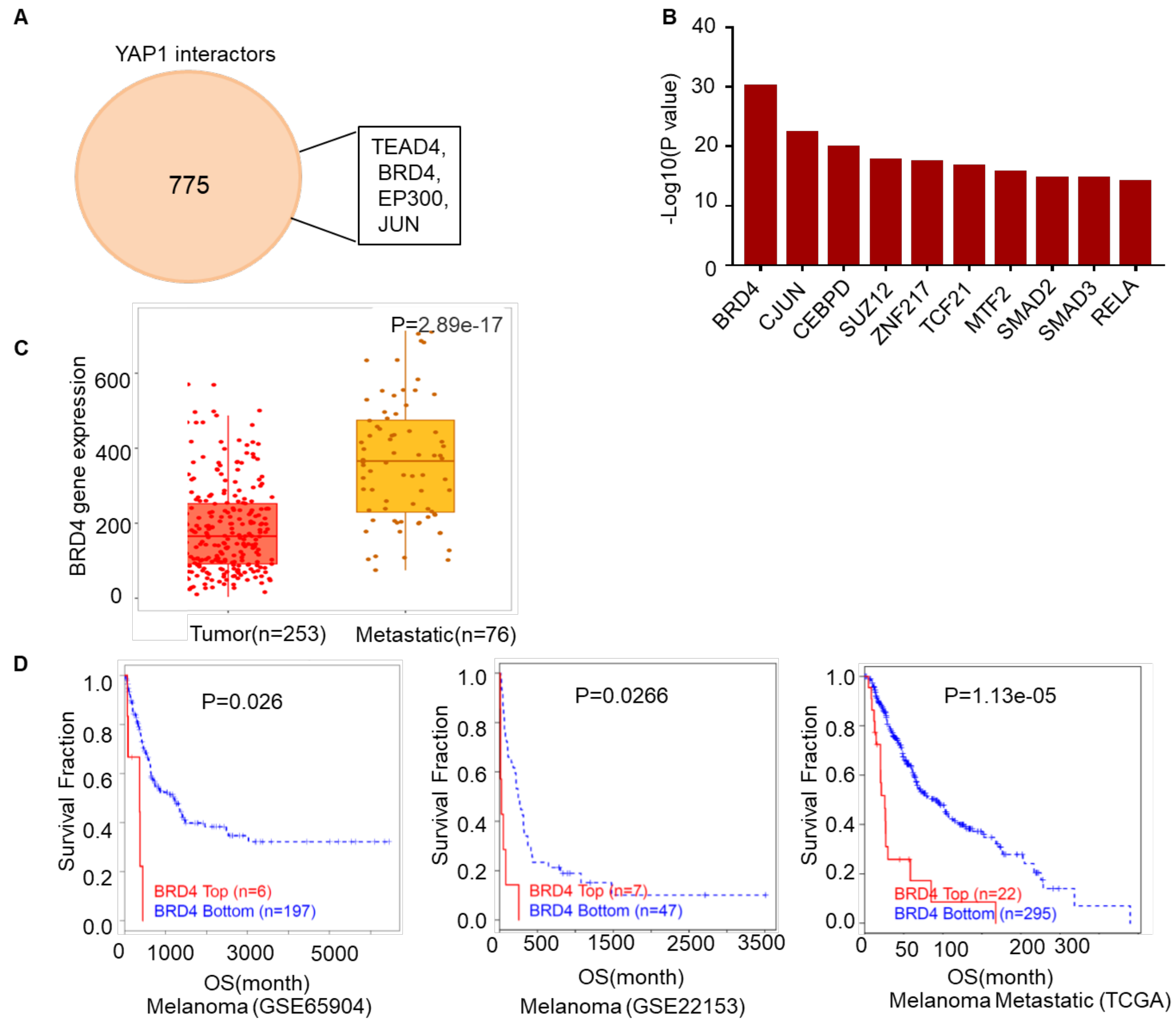


**Figure S4. BRAFi/MEKi resistant melanoma cells are sensitive to YAP co-effectors inhibitors.**

A) Biograd analysis of co-interactors YAP1. B) ChIP enrichment Analysis (ChEA) to analyze the transcription factors connected based on shared overlapping targets and binding site proximity in invasive and MEK inhibitor resistant melanoma. C) The expression of BRD4 in primary and metastatic melanoma patients. D) The overall survival curves comparing high- and low-expression levels of BRD4 in different SKCM cohorts (GSE65904; GSE22153; TCGA).

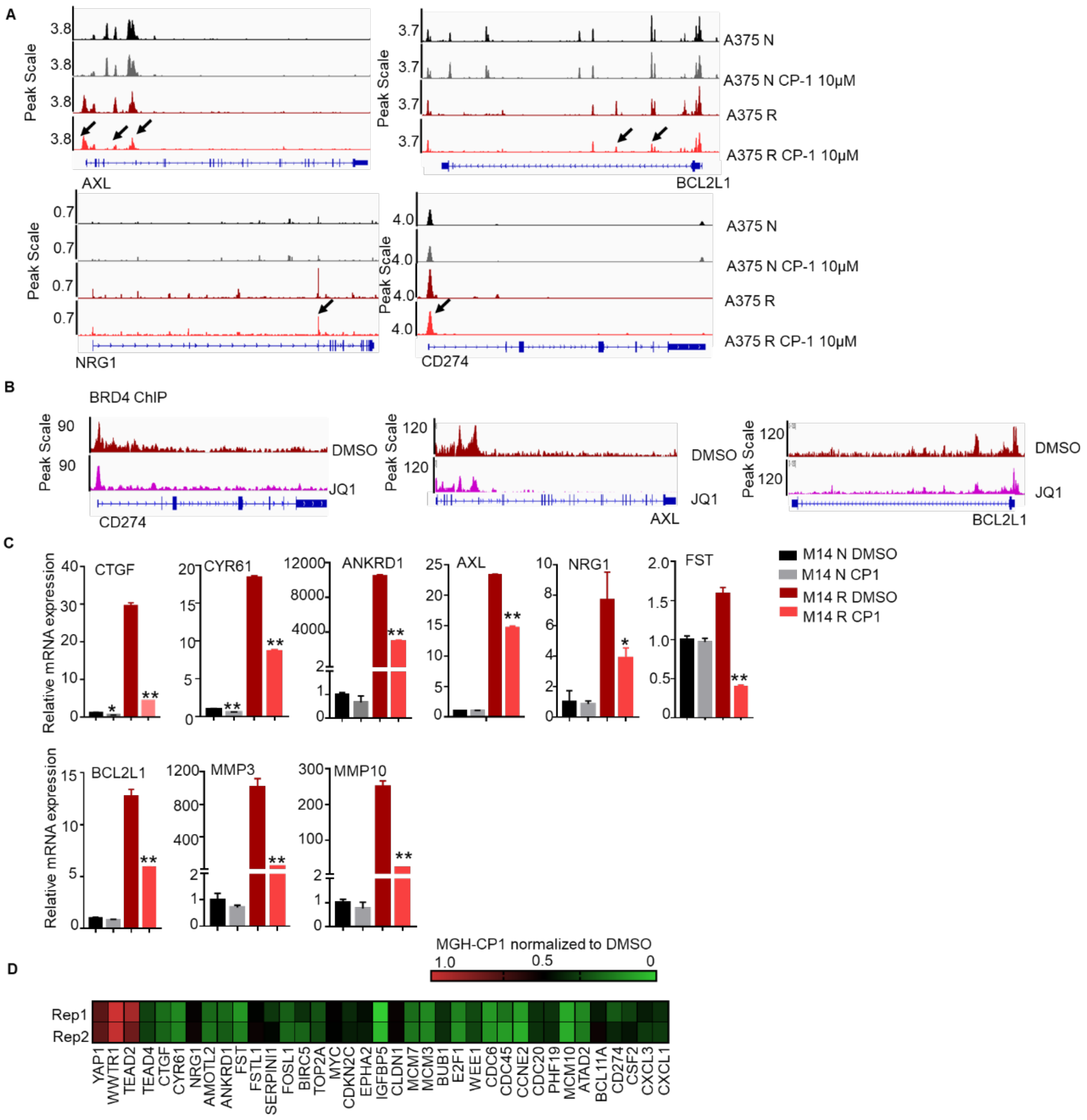


**Figure S5. Chromatin accessibility alterations in MEKi-Resistant compared with MEKi-sensitive melanoma cells after treatment by TEAD or BRD4 inhibitors.** A) ATAC-Sequencing analysis identifies alterations in chromatin accessibility regions in MEK inhibitor-resistant melanoma cells compared with MEK inhibitor-sensitive melanoma cells after treatment by MGH-CP1. B) The binding profiles of BRD4 in melanoma cells are depicted, with each track representing the normalized average occupancy for BRD4 in the promoter regions of *CD274*, *AXL*, and *BCL2L1* upon treatment with DMSO or JQ1. C) The relative mRNA expression level of YAP1 target genes was analyzed by qPCR after treatment with CP-1 in M14 Native and M14 BRAFi Resistant cells. The y axis shows the fold change in transcript levels versus DMSO-treated cells. Data are presented as mean ± S.D (n=3, *$P$<0.05, **$P$<0.01). D) RNA-seq dataset of MGH-CP1 in MDA-MB-231 cells (GSE177052). The heatmap illustrates the expression levels of Hippo pathway components and YAP1 targets after normalization to DMSO.

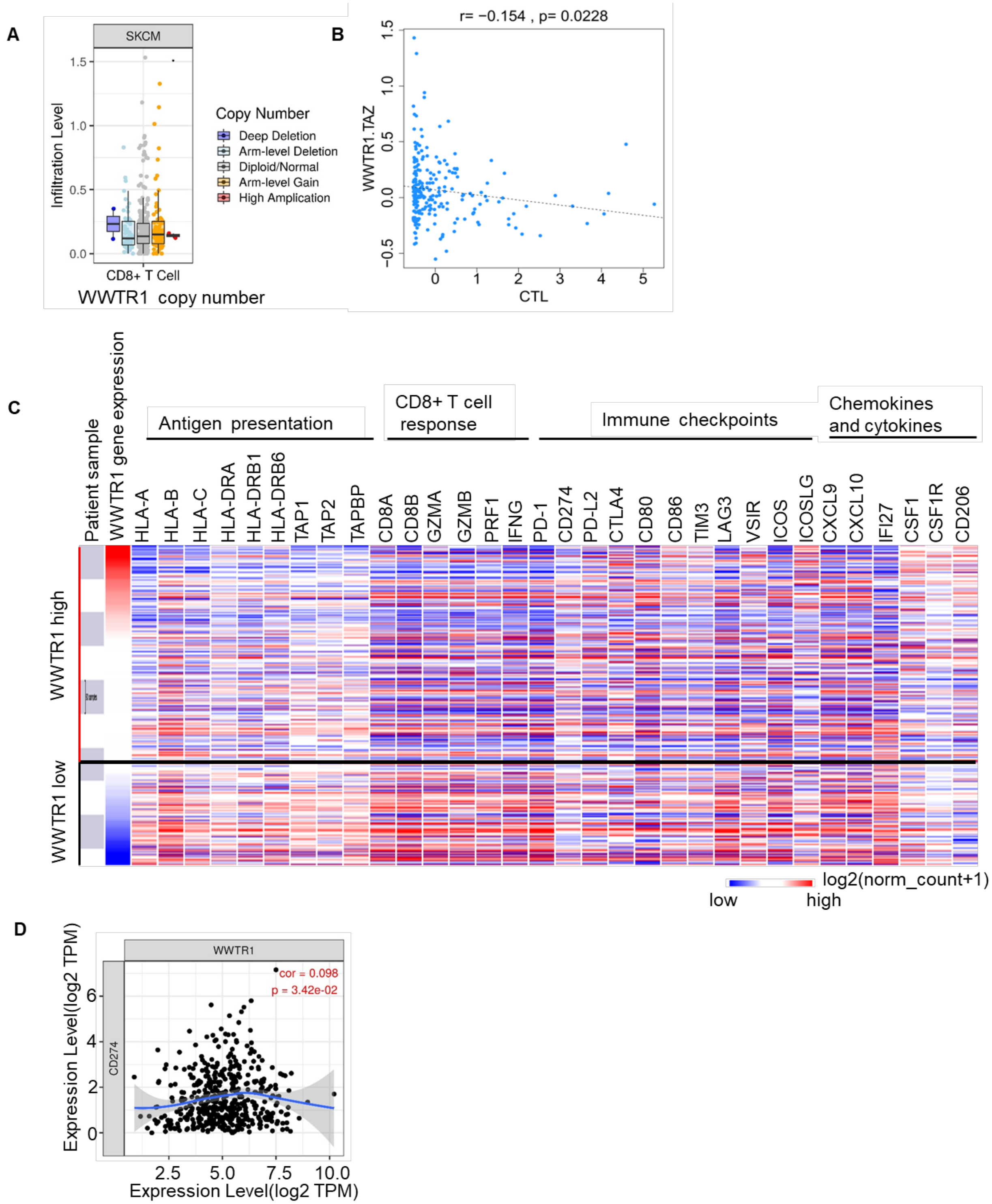


**Figure S6. The immune correlation of Hippo pathways component in melanoma patients' sample.** A, B) Analysis the association of copy number alteration of WWTR1 and WWTR1 protein expression with CD8+T or CTL (cytotoxic T cells) infiltration status. C) Xena Browser Visual Spreadsheet displays WWTR1, antigen presentation genes, CD8+ T cell signature genes, immune checkpoint, cytokines, and chemokines in melanoma patients. The expression is colored

red to blue for high to low expression. D) Analysis of the correlation between *WWTR1* and *CD274* in metastatic melanoma.

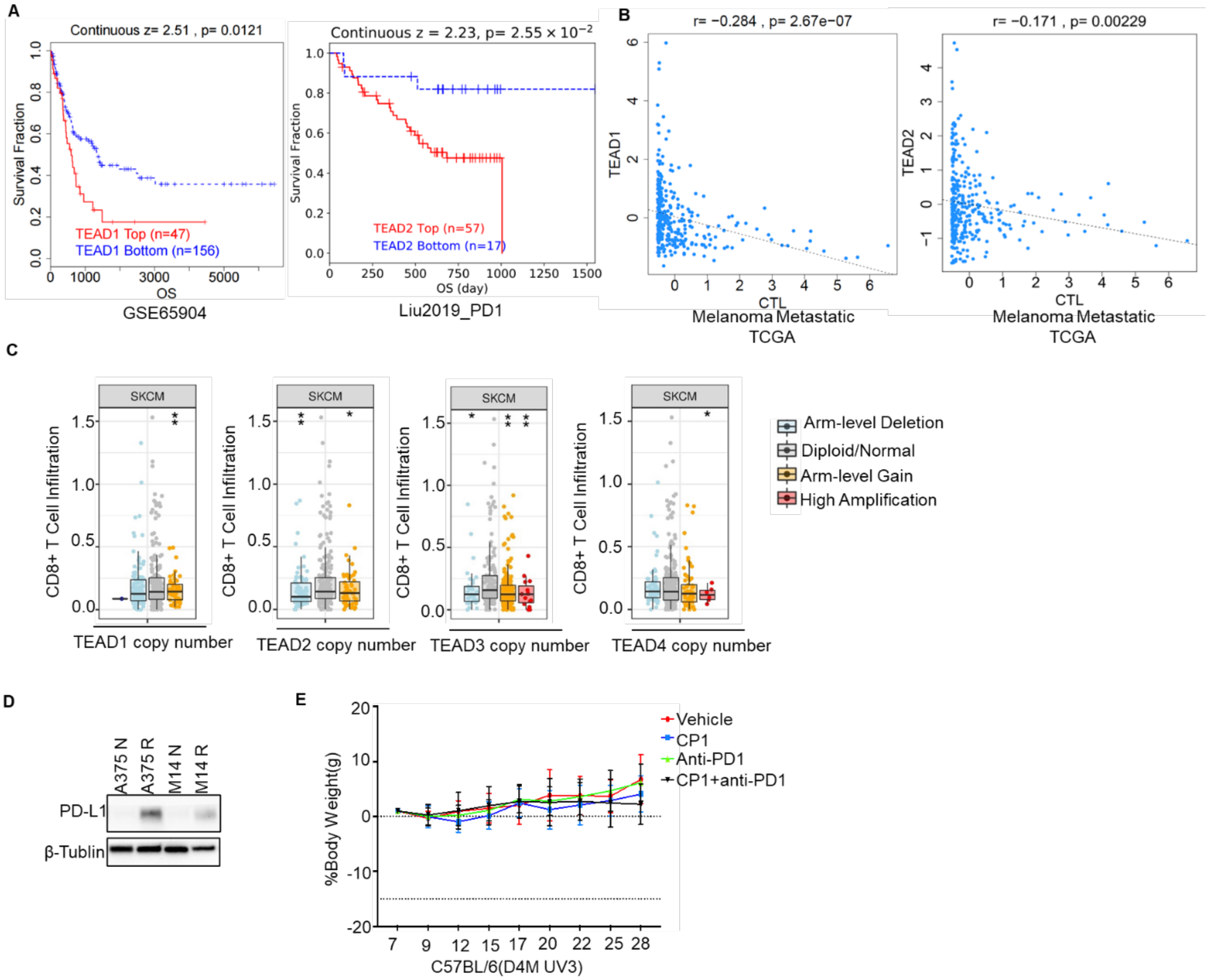


**Figure S7. TEAD inhibitor exerts an anti-tumor effect.** A) Kaplan-Meier survival curves showing the relationship between TEAD1 and TEAD2 expression and overall survival (OS) in melanoma. B) Analysis the association of TEAD1 and TEAD2 expression with CTL infiltration status. C) The association analysis of copy number alteration of TEAD1, TEAD2, TEAD3 and TEAD4 with CD8+T infiltration. D) Western blot analysis showing PD-L1 expression in melanoma cell lines. Top panel: PD-L1 levels in A375 (N: naïve, R: MEKi resistant) and M14 (N: naïve, R: BRAFi resistant) cells. Bottom panel: β-Tubulin as a loading control. E) Mice body weight was measured after treatment by the inhibitors or antibody in each group (n=7-8).

**Table S1.** List of primer sequences used for RT-qPCR analysis.

| qPCR Oligos | Sequence |
|---|---|
| YAP1 F | GCACCTCTGTGTTTTAAGGGTCT |
| YAP1 R | CAACTTTTGCCCTCCTCCAA |
| TAZ (WWTR1) F | GGCTGGGAGATGACCTTCAC |
| TAZ (WWTR1) R | CTGAGTGGGGTGGTTCTGCT |
| CTGF F | AGGAGTGGGTGTGTGACGA |
| CTGF R | CCAGGCAGTTGGCTCTAATC |
| CYR61 F | CCTTGTGGACAGCCAGTGTA |
| CYR61 R | ACTTGGGCCGGTATTTCTTC |
| ANKRD1 F | AGTAGAGGAACTGGTCACTGG |
| ANKRD1 R | TGGGCTAGAAGTGTCTTCAGAT |
| SERPINE1 F | TGGTGCTGATCTCATCCTTG |
| SERPINE1 R | AGAAACCCAGCAGCAGATTC |
| FSTL1 F | GCAGCAACTACAGTGAAATCC |
| FSTL1 R | ATGGCAGTTTCATTCTGTTCC |
| AXL F | CACCAGCAAGAGCGATGTGT |
| AXL R | CGGTCCTGGGGATTTAGCTC |
| FOSL1 F | AGCTGCAGAAGCAGAAGGAG |
| FOSL1 R | GGAGTTAGGGAGGGTGTGGT |
| NRG1 F | AACCTCAAGAAGGAGGTCAGC |
| NRG1 R | CTGGTTTCACACCGAAGGAC |
| H-EGFR-F | TGCACCTACGGATGCACTG |
| H-EGFR-R | CGATGGACGGGATCTTAGGC |
| H-EPHA2_F | GGAGGGATCTGGCAACTTGG |
| H-EPHA2_R | CTTCCTCCTGCGGTGGATAA |
| H BCL2L1 F | CTGAATCGGAGATGGAGACC |
| H BCL2L1 R | TGGGATGTCAGGTCACTGAA |
| BIRC3 F | GGAAATCCCCGAGTGGGTTT |
| BIRC3 R | CCAGTGGTTTGCATGTGCAC |
| MMP1 F | GACAGAAAGAGACAGGAGAC |
| MMP1 R | GAGTTATCCCTTGCCTATCC |
| MMP3 F | GCAGTTTGCTCAGCCTATCC |
| MMP3 R | GAGTGTCGGAGTCCAGCTTC |
| MMP10 F | GGCTCTTTCACTCAGCCAAC |
| MMP10 R | TCCCGAAGGAACAGATTTTG |
| MMP12 F | ACACATTTCGCCTCTCTGCT |
| MMP12 R | CCTTCAGCCAGAAGAACCTG |
| MITF-F | CATTGTTATGCTGGAAATGCTAGAA |
| MITF-R | GGCTTGCTGTATGTGGTACTTGG |
| SOX10-F | CCAGTACCCGCACCTGCAC |

| | |
|---|---|
| SOX10-R | CTTTCGTTCAGCAGCCTCCAG |
| TYR-F | TGCACAGAGAGACGACTCTTG |
| TYR-R | GAGCTGATGGTATGCTTTGCTAA |
| PMEL-F | AGGTGCCTTTCTCCGTGAG |
| PMEL-R | AGCTTCAGCCAGATAGCCACT |
| DCT-F | AACTGCGAGCGGAAGAAACC |
| DCT-R | CGTAGTCGGGGTGTACTCTCT |
| IL24-F | GCCTCTCAAATGCAGATGGT |
| IL24-R | GTCTTTCACAGCCCAGAAGG |
| CD74-F | GAGCTGTCGGGAAGATCAGA |
| CD74-R | AGGAAGTAGGCGGTGGTG |
| HLA-DRA-F | GCCAACCTGGAAATCATGACA |
| HLA-DRA-R | AGGGCTGTTCGTGAGCACA |
| CD274 F | ATGGTGGTGCCGACTACAA |
| CD274 R | TCCAGATGACTTCGGCCTT |
| STAT1-F | TGCAAAACCTTGCAGAACAG |
| STAT1-R | GGGCATTCTGGGTAAGTTCA |
| CXCL10 F | GTGGCATTCAAGGAGTACCTC |
| CXCL10 R | TGATGGCCTTCGATTCTGGATT |
| IGFBP5-F | TGACCGCAAAGGATTCTACAAG |
| IGFBP5-R | CGTCAACGTACTCCATGCCT |
| H FST1F | AAGACCGAACTGAGCAAGGA |
| H FST1R | TTTTTCCCAGGTCCACAGTC |
| CLDN1 F | GCATGAAGTGTATGAAGTGCTTGGA |
| CLDN1 R | CGATTCTATTGCCATACCATGCTG |
| 18s F | GTAACCCGTTGAACCCCATT |
| 18s R | CCATCCAATCGGTAGTAGCG |
| **ChIP-qPCR Oligos** | **Sequence** |
| H-NRG1 TSS F | GATTCAGTCCTGTGCTACGG |
| H-NRG1 TSS R | GCAGAGCTGATTCCTGACAC |
| HChIP-AXL F | TGAGTAGGGACCAGGGTTGG |
| HChIP-AXL R | CCACCACACAGACATGCACA |
| H-PDL1 ChIP-F | TCGGTCTGTGAAGGACTGC |
| H-PDL1 ChIP-R | ACCGTTGAGGAATGGATGAA |
| H BCL2L1 ChIP-F | GCAAAATGAGAGAGGGGGTGG |
| H BCL2L1 ChIP-R | GCAGAAAAGCATCAAGCTGTGG |
